# Glycoproteomic landscape and structural dynamics of TIM family immune checkpoints enabled by mucinase SmE

**DOI:** 10.1101/2023.02.01.526488

**Authors:** Joann Chongsaritsinsuk, Alexandra D. Steigmeyer, Keira E. Mahoney, Mia A. Rosenfeld, Taryn M. Lucas, Deniz Ince, Fiona L. Kearns, Alexandria S. Battison, Marie A. Hollenhorst, D. Judy Shon, Katherine H. Tiemeyer, Victor Attah, Catherine Kwon, Carolyn R. Bertozzi, Michael J. Ferracane, Rommie E. Amaro, Stacy A. Malaker

**Affiliations:** Department of Chemistry, Yale University, New Haven, CT 06511, USA; Department of Chemistry and Biochemistry, University of California, San Diego, La Jolla, CA 92093, USA; Department of Chemistry and Sarafan ChEM-H, Stanford University, Stanford, CA 94305, USA; Department of Pathology, Stanford University, Stanford, CA 94305, USA; Department of Medicine, Division of Hematology, Stanford University, Stanford, CA 94305, USA; Howard Hughes Medical Institute, Stanford University, Stanford, CA 94305, USA; Department of Chemistry, University of Redlands, Redlands, CA 92373, USA; Glycobiology Research and Training Center, University of California, San Diego, La Jolla, CA 92093, USA

## Abstract

Mucin-domain glycoproteins are densely O-glycosylated and play critical roles in a host of biological functions. In particular, the T cell immunoglobulin and mucin-domain containing family of proteins (TIM-1, −3, −4) decorate immune cells and act as key checkpoint inhibitors in cancer. However, their dense O-glycosylation remains enigmatic both in terms of glycoproteomic landscape and structural dynamics, primarily due to the challenges associated with studying mucin domains. Here, we present a mucinase (SmE) and demonstrate its ability to selectively cleave along the mucin glycoprotein backbone, similar to others of its kind. Unlike other mucinases, though, SmE harbors the unique ability to cleave at residues bearing extremely complex glycans which enabled improved mass spectrometric analysis of several mucins, including the entire TIM family. With this information in-hand, we performed molecular dynamics (MD) simulations of TIM-3 and −4 to demonstrate how glycosylation affects structural features of these proteins. Overall, we present a powerful workflow to better understand the detailed molecular structures of the mucinome.

## Introduction

Mucin-domain glycoproteins are characterized by extremely dense O-glycosylation that contributes to a unique, bottle-brush secondary structure which can extend away from the cell surface or form extracellular gel-like secretions.^1–3^ Mucin-type O-glycans are characterized by an initiating α-*N*-acetylgalactosamine (GalNAc) that can be further elaborated into several core structures which can contain sialic acid, fucose, and/or ABO blood group antigens.^4,5^ As a result, mucin domains serve as highly heterogeneous stretches of glycosylation that exert both biophysical and biochemical influence on the cellular milieu.^6,7^ The canonical family of mucins, e.g. MUC2 and MUC16, bear massive mucin domains that can reach 5-10 MDa in size and are heavily implicated in various diseases.^8,9^ That said, many other proteins contain mucin domains that do not necessarily reach that size or complexity. Indeed, we recently introduced the human “mucinome”, which comprises hundreds of proteins thought to contain the dense O-glycosylation that is characteristic of mucin domains.^10^ For instance, platelet glycoprotein 1bα (GP1bα) interacts with Von Willebrand Factor to mediate platelet adhesion, and mutations in GP1bα are involved in platelet-type Von Willebrand disease.^11^ C1 esterase inhibitor (C1-Inh) is a serine protease inhibitor and its deficiency is associated with hereditary angioedema.^12^ Finally, the T cell immunoglobulin and mucin domain containing protein family (TIM-1, TIM-3 and TIM-4) are critical regulators of immune responses and are highly implicated in various cancers.^13,14^ Though considerable progress has been made in the biological and analytical analyses of these and other mucin-domain glycoproteins, much remains unknown regarding their glycan structures, glycosylation site-specificity, and functional roles within the cellular environment.

This gap in knowledge is due, in part, to the challenges associated with studying mucins by mass spectrometry (MS).^15,16^ A typical MS workflow involves digesting proteins with workhorse proteases like trypsin, subjecting the peptides to separation via reverse phase HPLC, then analyzing them by high-resolution MS.^17^ Mucins present unique challenges at each stage of this process, but one of the most well-documented issues is the resistance of densely O-glycosylated domains to trypsin digestion.^3,5,18^ To address this challenge, several proteases have been introduced that selectively cleave at or near O-glycosylated residues thereby revolutionizing the field of O-glycoproteomics.^19–22^ These enzymes are aptly named O-glycoproteases; those that prefer mucin-domain glycoproteins are often termed mucinases. The first of these enzymes, OgpA (Genovis OpeRATOR), was characterized as an O-glycoprotease that cleaves N-terminally to glycosylated Ser or Thr residues but is hindered by the presence of sialic acid.^21,23^ Shortly thereafter, we introduced StcE as a mucinase that selectively digests mucin domains with a cleavage motif of T/S*_X_T/S, wherein the asterisk indicates a mandatory glycosylation site; we followed this work with a mucinase toolkit displaying a wide range of cleavage specificities.^19,20^ More recently, ImpA (NEB O-glycoprotease) was commercialized and, like OgpA, cleaves N-terminally to glycosylated Ser or Thr residues but, unlike OgpA, is less restricted by the glycans present.^24^ While these enzymes have aided in the analysis of many O-glycoproteins and mucins, each enzyme is accompanied by drawbacks: OgpA is limited by its resistance to sialic acid; the StcE cleavage motif is relatively restrictive; ImpA demonstrates preference for small amino acids adjacent to cleavage. Thus, an ideal, broad-specificity O-glycoprotease conducive to MS has not yet been characterized.

Another issue surrounding characterization of mucin-domain glycoproteins is that typical structural biology techniques are not well suited for glycoproteins, let alone the dense glycosylation characteristic of mucin domains. As covered in our recent review, current knowledge regarding mucin secondary structure originates from various low-resolution images generated by atomic force microscopy (AFM), scanning electron microscopy (SEM), and cryogenic electron microscopy (cryoEM)^3,25–27^ While this has allowed us to definitively visualize the linearity of the mucin protein backbones, by nature of the techniques, we are unable to (a) discern the individual glycans and how they contribute to changes in protein structure, or (b) observe the mucin protein dynamics. Additionally, these methods often require large (>50 kDa), pure, and concentrated protein samples, which are difficult to obtain with most native mucin-domain glycoproteins.^3^ More recently, and eloquently reviewed in reference 28, many advances have been made in computational modeling of glycoproteins.^28^ These molecular dynamics (MD) simulations have revealed some of the many roles that glycans play in the structure, stability, dynamics, and function of glycoproteins. Most notably, Amaro and colleagues revealed, for the first time, the functional role of the glycan shield in the activation mechanism of the SARS-CoV-2 spike protein.^29,30^ That said, while mucins have been subjected to MD simulations previously, they are often modeled with a static glycan structure, lack precise glycosylation information, and are simulated with coarse grained MD. Taken together, for mucin-domain glycoproteins, we generally do not know the glycoproteomic landscape nor how the glycans work in concert to control protein and cellular dynamics.

Here, we present a powerful technique to map the complex glycosylation within mucin domains and pair this information with MD simulations in order to better understand how glycans affect glycoprotein secondary structure and dynamics. We first introduce a new mucinase, *Serratia marcescens* Enhancin (SmE), and demonstrate that its unique ability to cleave at glycosites decorated by a myriad of glycans enabled enhanced glycosite and glycoform analysis by MS. We next showed that SmE outperforms the commercial glycoproteases OgpA and ImpA for these purposes, and we used molecular modeling to understand its uniquely broad tolerance for dense glycosylation. With SmE in-hand, we then obtained complete O-glycoproteomic information for all TIM family proteins and demonstrated that TIM-3 has markedly fewer O-glycosites when compared to TIM-1 and −4. To better understand how these glycans affect overall protein structure, we then employed MD simulations and showed that TIM-3 has a much shorter persistence length and higher flexibility than TIM-4, primarily attributed to the dense glycosylation in the latter. Overall, this workflow aids in unraveling the complex molecular mechanisms behind mucin domains, their glycan patterns, and their contribution to cellular biology.

## Results

### Characterization of SmE cleavage motif and glycan specificity

Various microorganisms found within mucosal environments secrete proteolytic enzymes that have been shown to be advantageous tools for MS analysis of mucins. We and others have mined the microbiota to generate a toolkit of O-glycoproteases, each having unique peptide and glycan specificities.^19,20,22,24^ In particular, *Serratia marcescens* is a pervasive opportunistic pathogen in humans. This organism secretes a mucinase, SmE, which is a viral enhancin protein shown to promote arboviral infection of mosquitoes by degrading gut membrane-bound mucins.^31^ Like previously characterized O-glycoproteases BT4244 and ImpA, SmE contains a catalytic domain belonging to the Pfam family PF13402 (peptidase M60, enhancin, and enhancin-like or M60-like family) that is defined by a conserved HEXXH metallopeptidase motif.^32^ We expressed SmE as a 94-kDa soluble, N-terminal His-tagged protein in *E. coli* at a high-yield expression level of 65 mg/L (**Figure S1**). To determine SmE’s mucin selectivity, we digested glycoproteins with and without mucin domains at a 1:20 enzyme to substrate (E:S) ratio. SmE preferentially cleaved the mucin proteins C1 esterase inhibitor (C1-Inh), CD43, and TIM-1, whereas it did not significantly cleave the non-mucin glycoproteins fibronectin and fetuin (**Figure S2**).

As in our previous work, we then characterized SmE’s cleavage motif using biologically relevant mucin-domain glycoproteins (C1-Inh, TIM-1, TIM-3, TIM-4, and GP1bα). These proteins were digested with SmE and subjected to MS analysis (**Figure 1A**). Manually validated glycopeptides were mapped to protein sequences to identify sequence windows, which were input into weblogo.berkeley.edu to determine minimum sequence motifs. As demonstrated in **Figure 1B**, SmE cleaved N-terminally to a glycosylated Ser or Thr. SmE accommodated a variety of O-linked glycans at the P1’ position (pie chart, right), including sialylated core 1 and core 2 O-linked glycans; surprisingly, it also tolerated fucosylated ABO blood group antigens. SmE was also able to accommodate O-glycosylation at the P1 position (pie chart, left). Given the higher percentage of smaller O-glycan structures (GalNAc, GalNAc-Gal), it appears that the P1 O-glycosylation tolerance was less permissive at this position. However, these pie charts represent only site-localized glycan structures; site-localization at the C-terminus of the peptide is more challenging due to a lack of positive charge. Thus, the apparent preference for smaller glycan structures at the P1 position is likely due to issues in glycoproteomic analysis as opposed to an inability for SmE to cleave at residues bearing larger glycans.

**Figure 1.**
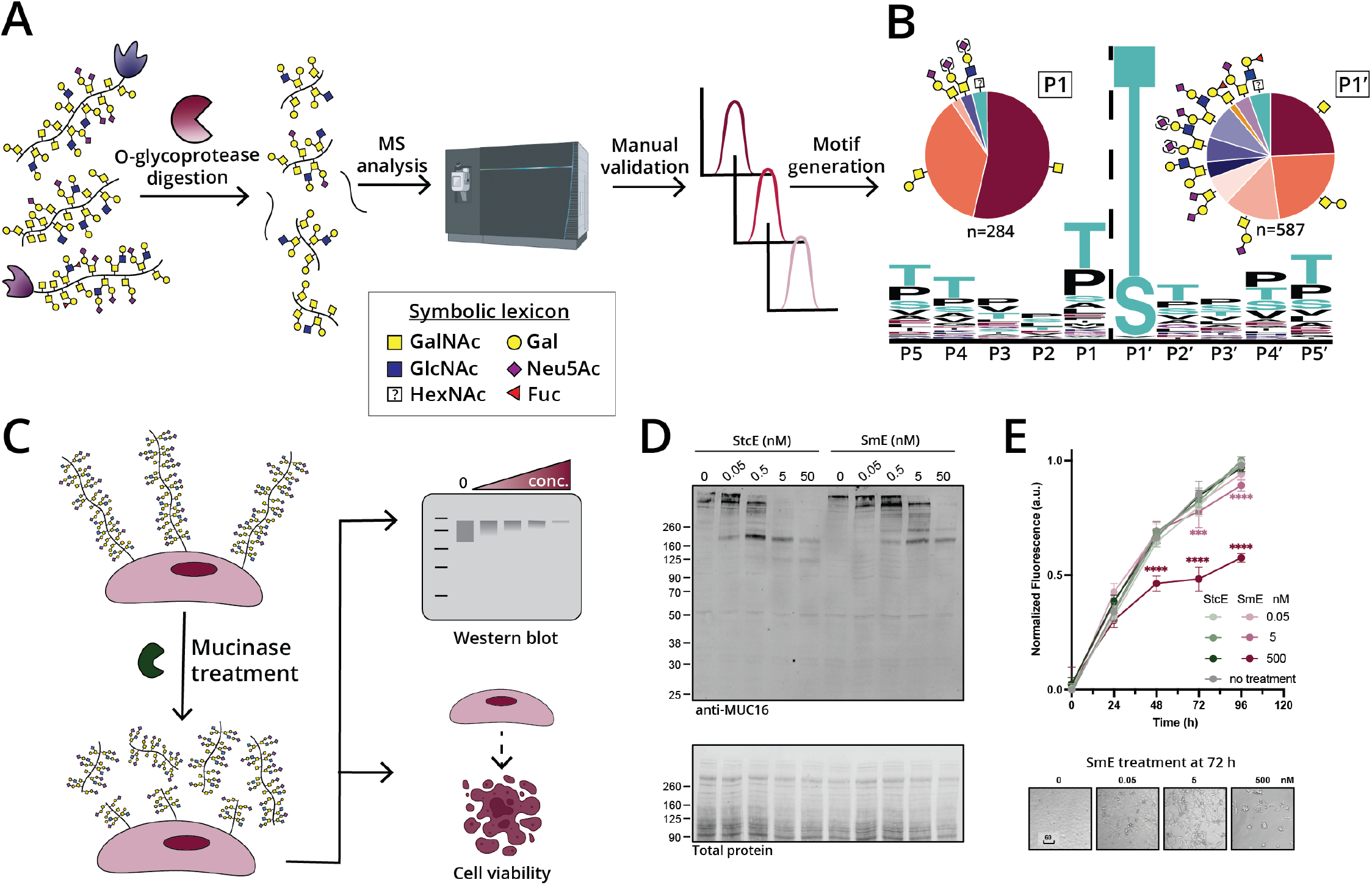
Characterization of mucinase SmE for analysis and degradation of mucin-domain glycoproteins. (A) Workflow for generating consensus sequence of SmE. (B) Five recombinant mucin-domain glycoproteins were digested with SmE and subjected to MS analysis. Peptides present in the mucinase-treated samples were used as input for weblogo.berkeley.edu (±5 residues from the site of cleavage). Parentheses around sialic acids (purple diamond) indicate that its linkage site was ambiguous. (C) Workflow to evaluate the toxicity and cell surface activity of SmE. (D) HeLa cells were treated with StcE (left) or SmE (right) at the noted concentrations for 60 min. Following treatment, the cells were lysed in 1X NuPAGE LDS Sample Buffer with 25 mM DTT, subjected to separation by gel electrophoresis, and probed for MUC16 via Western blot. Proteins were transferred to a nitrocellulose membrane using the Trans-Blot Turbo Transfer System (Bio-Rad) at a constant 2.5 A for 15 min. Total protein was quantified using REVERT stain before primary antibody incubation overnight at 4 °C. An IR800 dye-labeled secondary antibody was used according to manufacturer’s instructions for visualization on a LiCOR Odyssey instrument. (E) HeLa cells were treated with SmE and StcE at 0, 0.05, 5, and 500 nM. At t = 0, 24, 48, 72, 96 hours post treatment, PrestoBlue was added according to manufacturer’s instructions. After 2 hours, the supernatant was transferred to a black 96 well plate and analyzed on a SPECTRAmax GEMINI spectrofluorometer using an excitation wavelength of 544 nm and an emission wavelength of 585 nm. Statistical significance was determined using the two-way ANOVA analysis in Graphpad PRISM software and is reported with respect to the no mucinase control condition. *** p <0.001, **** p <0.0001. Scale bar = 60 μm.

SmE provides a complementary cleavage profile to StcE which, as mentioned above, cleaves at a T/S*_X_T/S motif. In that work, we also demonstrated that StcE is non-toxic to cells and can be employed to release mucins from the cell surface.^19^ Many researchers have since used our mucinase toolkit, especially StcE, to remove mucins from the cell surface and/or degrade mucins in biological samples.^1,33–36^ Importantly, the MUC1 repeat sequence HGVTSAPDTRPAPGSTAPPA does not contain StcE’s cleavage motif, so limited digestion occurs within this region.^37^ Given that SmE has a complementary cleavage motif, a combinatorial treatment strategy could enable further degradation of mucins from various biological samples. Thus, we sought to understand whether SmE is similarly capable of digesting mucins from the cell surface, and whether the enzyme is likewise non-toxic to cells (**Figure 1C**). Treating HeLa cells with SmE resulted in a reduction in MUC16 staining by Western blot in a manner comparable to that of StcE (**Figure 1D**). We also detected released MUC16 fragments in the supernatant of SmE treated cells (**Figure S3**). Importantly, SmE is not toxic to cells under conditions used previously for StcE, although cell death is observed at higher SmE concentrations over longer treatment durations (**Figure 1E** and **Figure S4**). Taken together, these results indicate that SmE, like StcE, can effectively cleave mucins from the cell surface as a tool to probe mucin biological function.

In summary, SmE is a mucinase with the unique ability to cleave between two glycosylated residues bearing complex O-linked glycans. We anticipate that the characteristics of SmE’s cleavage motif and glycan specificities will facilitate not only improved glycoproteomic mapping of mucin-domain glycoproteins, but also clearance of mucins from biological samples.

### SmE outperforms commercial O-glycoproteases OgpA and ImpA for mucin analysis

To compare the activity of SmE in context with widely used, commercially available O-glycoproteases, we decided to benchmark against both OgpA and ImpA.^21,24,38,39^ OgpA was originally identified in *Akkermansia muciniphila*, a commensal bacterium known to regulate mucin barriers through controlled degradation.^40^ As mentioned above, OgpA is reported to cleave N-terminally to O-glycosylated Ser or Thr residues, with highest affinity towards asialylated core 1 species.^41,42^ Thus, typical workflows with this enzyme involve removal of sialic acids, which limits its use to site-mapping as opposed to providing information on native glycan structures. ImpA is derived from *Pseudomonas aeruginosa*, an opportunistic bacterial pathogen that can cause severe infection.^43^ Like OgpA, ImpA cleaves N-terminally to an O-glycosylated Ser or Thr residue; however, this enzyme has been reported to accommodate more complex, sialylated glycans, expanding its glycoproteomic potential beyond that of OgpA. Efficiency of cleavage by ImpA has been observed to be influenced by amino acid identity in the P1 position. For example, reduced efficiency was seen when P1 was occupied by Arg or Ile, and cleavage was not observed when occupied by Asp.^24,39^

To directly compare the activities of OgpA, ImpA, and SmE, mucin-domain glycoproteins TIM-1, −3, −4, GP1bα, and C1-Inh were digested in the presence and absence of sialidase, followed by gel electrophoresis (**Figure S5**), MS analysis, and manual glycopeptide validation (see **SI Tables 1-5** for all annotated glycopeptides). Additionally, we included fetuin to investigate the enzymes’ selectivity for mucin glycoproteins (**SI Table 6**). As demonstrated in **Figure 2A-B**, we confirmed the reported cleavage motifs and glycan preferences of both OgpA and ImpA. Notably, after ImpA digestion, we did not detect any glycosites in the P1 position, suggesting that ImpA does not cleave between two glycosylated residues. Given that mucin domains contain many neighboring O-glycosites, this presents a significant limitation in the use of ImpA for mucinomic analysis.

**Figure 2.**
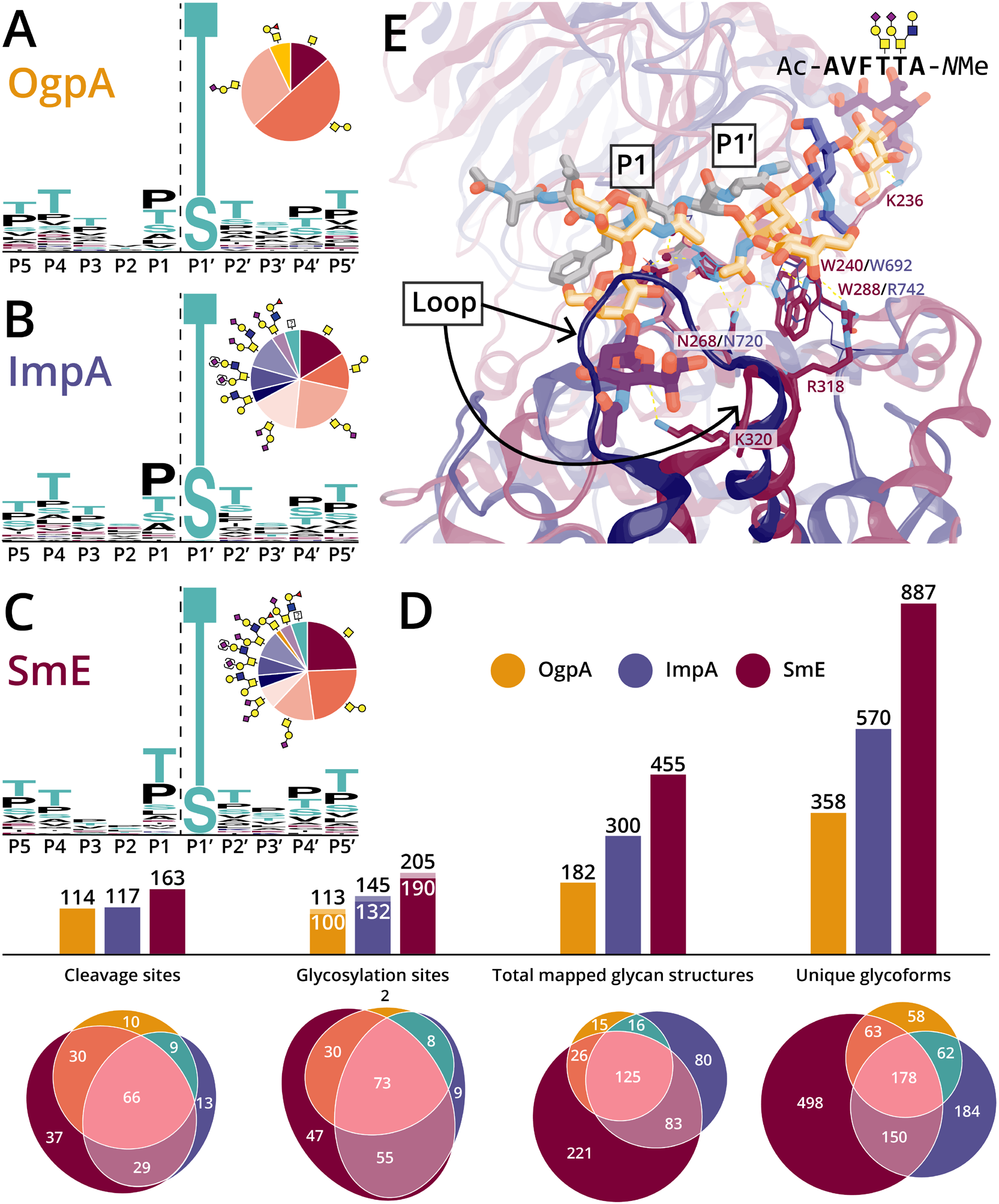
SmE outperformed commercial O-glycoproteases due to its structural permissiveness. Cleavage motifs for (A) OgpA, (B) ImpA, and (C) SmE as determined by digestion followed by MS and manual curation of glycopeptides. (D) Bar graphs and Euler plots demonstrating counts and overlap between enzymes regarding the number of observed cleavage sites, localized glycosylation sites, total glycan structures, and unique glycoforms. For the “Glycosylation sites” bar graph, glycosites localized via MS are denoted by white numbers; black numbers above include implied glycosites where cleavage was observed but the glycosite was not localized. (E) A glycopeptide docked in the active site of SmE (maroon) and ImpA (blue), highlighting differences between key loops and residues of the two O-glycoproteases.

In contrast, we found that SmE activity was not limited by glycan complexity or adjacent glycosylation (**Figure 1B, 2C**). Perhaps for this reason, digestion with SmE greatly improved the depth and coverage of the glycoproteomic landscape for each mucin-domain glycoprotein we investigated. Number of cleavage sites, unique O-glycosites, mapped glycan structures, and total glycoforms were determined (**Figure 2D**, see **Figure S6** for maps of all cleavage events). Note that for cleavage sites and unique O-glycosites, we included the sialidase treated samples; however, for the total mapped glycan structures and unique glycoforms, we only considered digests without sialidase treatment, as its use inherently limits identification of native glycan complexity. Here, total mapped glycan structures are calculated by counting every O-glycan associated with each O-glycosite. Unique glycoforms refers to the total number of validated glycopeptides that were identified from protein digestions. In our analyses, we found that OgpA allowed for the identification of 113 glycosites, 182 mapped glycan structures, and 358 unique glycoforms. ImpA demonstrated significant improvement over OgpA, enabling localization of 145 glycosites, 300 mapped structures, and 570 glycoforms. Even more impressively, SmE digestion allowed us to identify 205 glycosites, 455 mapped structures, and 887 glycoforms. Notably, use of SmE permitted the identification of 47 unique glycosites, 221 glycan structures, and 498 glycoforms that were not detected using the other enzymes. Previously, glycomic and glycoproteomic analyses of C1-Inh hinted at a total of approximately 25 O-glycosites; however, these were sparingly localized to individual residues.^12,44,45^ SmE enabled full glycoproteomic mapping of the C1-Inh mucin domain (**Figure S7, SI Table 1**), thus reinforcing the utility of this enzyme. Taken together, SmE greatly outperformed both OgpA and ImpA with regard to glycoproteomic analysis of mucin domains.

Although SmE clearly exhibited benefits for mucin glycoprotein analysis, we observed certain limitations associated with its use. Relative to its performance in mucin domains, the efficiency of SmE was greatly diminished when used on non-mucin glycoproteins, such as fetuin (**Figure S6; SI Table 6**). Analysis by MS revealed that SmE was able to cleave non-mucin glycoproteins to a limited extent; however, the abundance of fetuin glycopeptides resulting from SmE digestion was significantly reduced when compared to those of ImpA, and in some cases, OgpA (**Figure S8**). We also observed that SmE digestion was most effective against large (>50 residue) mucin domains. These observations, along with the digestion assay in **Figure S2**, support the notion that SmE is a mucin-selective O-glycoprotease. However, while SmE greatly enhanced sequence coverage and depth, some glycosylation was only localized through the use of OgpA and ImpA (**Figure 2D**). For this reason, we recommend a complementary, multi-enzyme approach to fully elucidate the glycoproteomic landscape of mucin-domain glycoproteins. Additionally, since ImpA was previously reported to have P1 selectivity, we generated “anti-logos” to determine surrounding residues that were unfavorable for cleavage. Here, we considered the total cleavage maps depicted in **Figure S6**, and whenever an enzyme did not cleave at an observed cleavage site, we took the surrounding amino acids and generated a logo. Interestingly, we found that SmE has a lower cleavage efficiency at Ser residues (**Figure S9**). While this could be because Ser is less often decorated with mucin-type O-glycosylation, it might be an important consideration for digestion of mucin domains bearing high levels of Ser residues. Finally, we observed that SmE exhibited reduced proteolytic activity on recombinant proteins expressed in murine-derived cell lines, as exemplified by (a) the digestion of TIM-1 from NS0 and HEK cells over the course of six hours (**Figure S10**) and (b) limited digestion of CD43 derived from NS0 cells (**Figure S2**).

### Molecular modeling helps rationalize different substrate selectivity between SmE and ImpA

Previously, we used molecular docking to better understand StcE’s substrate selectivity.^19^ Given that SmE and ImpA have catalytic domains belonging to the same Pfam family,^46,47^ yet have quite different cleavage motifs, we decided to again use molecular modeling to understand the structural basis behind these differences. In addition to its catalytic domain, SmE has two mucin-binding modules (PF03272) while ImpA has unique helical (PF18642) and N-terminal (PF18650) domains. OgpA, by contrast, is a single-domain enzyme with a catalytic metzincin motif that is not defined by Pfam and is more distantly related to SmE and ImpA, and therefore was excluded from these comparison analyses.

To date, four unique crystal structures of ImpA have been determined,^32,48^ including one with a ligand bound at the active site and a second with a ligand bound at an exosite located in the N-terminal domain (PF18650). The structure of SmE, on the other hand, has not yet been elucidated. As such, we aligned structures of all characterized enzymes with a PF13402 catalytic domain,^49–52^ including the SmE structure recently predicted by AlphaFold (**Figure S11**).^53,54^ We then docked a TIM-4-based bisglycosylated peptide into the predicted SmE structure to provide insight into potential substrate recognition.

In its cocrystal structure with (Gal-GalNAc)Ser,^32^ ImpA uses specific residues to recognize glycans branching from P1’ (**Figure 2E,** blue). The side chains of the conserved residues Trp692 and Asn720 form polar contacts with the carbonyl of the N-acetyl group of the GalNAc moiety, and the side chain of Arg742 also interacts with the 3-OH and 4-OH of GalNAc; the Gal moiety, on the other hand, does not interact with the enzyme and is projected into solvent. The helix lining the active site of ImpA is also short,^38^ indicating that branched glycans – though absent from the crystallized ligand - could be reasonably accommodated by this enzyme in a similar manner to ZmpB and ZmpC.^32^ These combined factors likely impart ImpA with activity on substrates bearing mucin-like glycosylation at P1’ but little selectivity for particular modifications beyond the initiating GalNAc moiety, consistent with our MS findings.

In our docked structure of SmE with a bisglycosylated TIM-4-based glycopeptide (**Figure 2E,** red), we found analogous interactions between conserved residues Trp240 and Asn268 and the GalNAc initiating from P1’. Unique contacts were found between the 3-OH of GalNAc and the side chain of Trp288, rather than with an Arg residue as seen in crystal structures of ImpA and other PF13402-containing enzymes. While there is an Arg residue (Arg282) nearby in the sequence, it is not predicted to flank the GalNAc moiety in the AlphaFold structure, and we found that this residue is less conserved in PF13402-containing enzymes than previously suggested (**Figure S11**).^49^ Interestingly, a different Arg side chain (Arg318) contacted the 4-OH of the Gal residue, likely imparting further specificity for mucin-like glycosylation. Similar to ImpA, SmE is predicted to have a short active site helix. Here, the branched glycan was well accommodated by the enzyme, and we commonly found orientations that formed contacts between the ligand and the active site helix as well as the preceding loop, similar to what was observed between ZmpB/ZmpC and their branched ligands (**Figure S12**).^32,50^ Together, these results suggest that SmE has better recognition of the initiating GalNAc and Gal residues, and that it likely forms additional interactions with branched glycans.

Neither the ImpA crystal structure nor the SmE predicted structure contain a beta hairpin analogous to the one found to recognize P1 glycans by the mucinase AM0627 (**Figure S11, S13**), which also allows glycosylated P1 Ser/Thr.^52^ Thus, the steric environment in this region is primarily defined by a single loop that is significantly larger in ImpA (Trp770-Leu778) than it is in both SmE (Asn313-Asp316) and AM0627 (Leu384-Asp388). In our docked structure, we observed that the short loop of SmE allowed the enzyme to easily accommodate the P1 glycan, with neither the GalNAc nor the Gal residue forming direct contacts with the enzyme; the sialic acid residue could interact with the enzyme or project toward solvent. This is in contrast to AM0627, which forms direct contacts between its beta hairpin and the GalNAc and Gal residues to impart requirement for P1 glycosylation.^20,49,52^ The long loop in ImpA, by contrast, sterically clashes with all three subunits of the P1 glycan, which explains why ImpA is unable to cleave between adjacent residues bearing glycosylation. More broadly, these and prior findings suggest a delicate interplay between the hairpin and loop in determining this enzyme family’s tolerance, preference, or requirement for particular glycans at P1 (**Figure S13**). In the case of SmE, the short loop and absence of a hairpin allows the enzyme to tolerate (but not require) larger glycans at the P1 position, which again supports our MS findings.

### Molecular modeling identifies potential secondary mucin binding in SmE

Multidomain mucinases are hypothesized to arrange their noncatalytic domains into an architecture that enables specific recognition of secondary sites along the linear bottle-brush of mucins,^50,55^ and recombinant StcE lacking one of these noncatalytic domains showed reduced activity on mucin substrates.^56,57^ Interestingly, the related metalloprotease MMP-1 (collagenase) required an accessory domain to bind and cleave the linear triple helix of collagen;^58^ structural and functional studies revealed the importance of cooperativity between this enzyme’s catalytic and accessory domains as well as specific interactions between each domain and the collagen triple helix.

In one crystal structure, ImpA binds a glycopeptide in an exosite located within its noncatalytic N-terminal domain (PF18650).^48^ During our initial docking study, we observed that one of the accessory mucin-binding modules (Asn537-Leu650, PF03272) in SmE is also positioned to recognize additional sites in mucin substrates. At present, there is no experimentally determined structure of a PF03272 domain; thus, the precise details of the domain’s structure and ligand recognition remain unknown. As such, we grafted larger segments of TIM-4 (*vide infra*) onto the docked glycopeptide to determine if this accessory domain can potentially bind the substrate. We observed that several of the larger TIM-4 substrates positioned glycans adjacent to a predicted binding pocket formed from two conserved segments in the mucin-binding module (**Figure S14, S15**). While additional work is required to validate this initial finding, the result suggests that SmE could use its mucin-binding module to cooperatively bind mucin substrates and sterically occlude more globular O-glycoproteins. Such a model would explain SmE’s observed preference for mucins over non-mucin O-glycoproteins like fetuin.

### Glycoproteomic mapping of TIM-1, −3, and −4

With this new tool in-hand, we reasoned that SmE could be used to sequence immune checkpoint mucin-domain glycoproteins at the molecular level. In particular, TIM-1, −3, and −4 are key players in immune cell function and are predicted to be modified by many O-glycosylation sites.^59^ Each protein contains an N-terminal variable immunoglobulin (IgV) domain followed by a densely glycosylated mucin domain of varying length, a single transmembrane domain, and a C-terminal intracellular tail.^60^ TIM-3 is highly implicated in cancer pathways, thus much of the literature to date has focused on better understanding its molecular interactions. In short, when TIM-3 is not bound to its extracellular ligands (including phosphatidylserine (PtdSer), high mobility group box 1 protein (HMGB1), and/or galectin-9 (Gal-9)) via its IgV domain, the TIM-3 cytoplasmic tail induces phosphorylation of the T cell receptor (TCR), which promotes T cell proliferation and survival. However, when TIM-3 is bound to one of its ligands, the cytoplasmic tail becomes phosphorylated and consequently promotes a state of T cell exhaustion that is characteristic of many cancers (**Figure 3A**, left).^61^ As such, several antibodies against TIM-3 are currently being investigated as cancer immunotherapies, often in combination with canonical checkpoint inhibitors like PD-1.^62–64^

**Figure 3.**
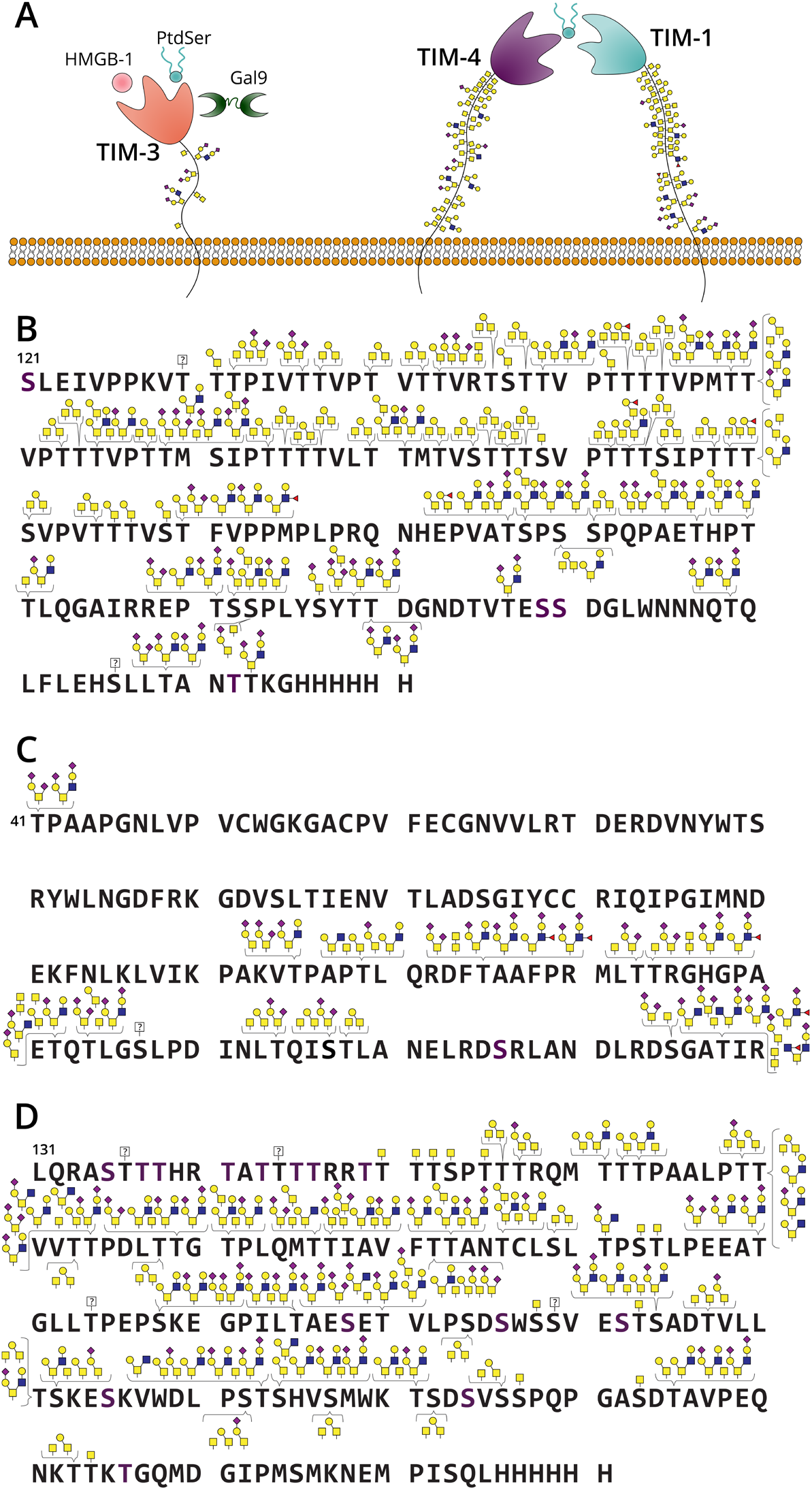
Glycoproteomic mapping of TIM family proteins. (A) Cartoon of TIM family structure and ligand interactions. TIM-3 interacts with ligands PtdSer, HMGB1, and/or Gal-9; through intracellular signaling these interactions deactivate T cell function and cytokine release. TIM-1 and TIM-4 purportedly interact through PtdSer to enact effector function. Recombinant TIM-1 (B), TIM-3 (C), and TIM-4 (D) were subjected to digestion with SmE, ImpA, OgpA, and/or trypsin followed by MS analysis and manual data interpretation. Brackets indicate glycans sequenced at each Ser/Thr residue at >5% relative abundance. For full glycoproteomic sequencing data, see SI Tables 4, 7, and 8.

Compared to TIM-3, little is known about TIM-1 and TIM-4, potentially because these proteins are predicted to bear more O-glycosites than TIM-3, thus complicating their analysis. However, it is known that the combination of TIM-1 blockage and TCR stimulation promotes T cell proliferation and cytokine production;^65^ TIM-4 is a PtdSer receptor and binds when PtdSer is exposed on the surface of apoptotic cells (**Figure 3A**, right).^66^ While much remains to be discovered about TIM-1 and −4, it is apparent that the entire TIM family plays critical roles in regulating immune responses in normal and dysregulated cellular states. However, only predicted glycosylation sites in the TIM family have been discussed in the literature, leaving their true glycoproteomic landscape a mystery. It follows, then, that we also do not understand how glycosylation contributes to TIM protein ligand binding, structural dynamics, and intracellular signaling. Ultimately, this lack of information hampers our understanding of the TIM family structure and function, which could have strong implications for cancer immunotherapy.

According to NetOGlyc 4.0,^67^ TIM-1, −3, and −4 were predicted to bear 67, 8, and 66 O-glycosites, respectively. Additionally, we recently developed a “Mucin Domain Candidacy Algorithm” which takes into account predicted O-glycosites, glycan density, and subcellular location in order to output a “Mucin Score”.^10^ This value was developed as a method to gauge the likelihood that a protein contains a mucin domain; a protein receiving a score above 2 indicated a high probability. Interestingly, TIM-1 and TIM-4 scored above 6, whereas TIM-3 received a score of 0.^10^ Beyond the biological implications of these proteins, we were curious to understand the glycoproteomic landscape of the TIM family given the large disparity in predicted O-glycosites and Mucin Scores. Thus, we digested the three recombinant TIM protein ectodomains with SmE and performed MS analysis followed by manual curation of the glycopeptides. Given our earlier observations regarding SmE’s resistance to proteins bearing sparse glycosylation, we digested TIM-3 with ImpA and OgpA to ensure full sequence coverage and O-glycosite identification. As seen in **Figure 3B** with a full list of annotated glycopeptides in **SI Tables 4, 7-8**, we identified all 67 of the predicted O-glycosites on TIM-1; many of these sites were modified by a myriad of O-glycans, thus demonstrating the massive microheterogeneity in mucin domains. TIM-4 was similarly dense in glycosylation and we site-localized a total of 51 O-glycosites (**Figure 3D**). In contrast, only 14 sites of O-glycosylation were detected on TIM-3; the glycosylation was also much less dense, though still quite heterogeneous (**Figure 3C**). Generally, recombinantly expressed proteins are thought to display relatively simple glycosylation (e.g., core 1 or 2 structures). Intriguingly, despite the fact that these proteins were recombinantly expressed in HEK293 cells, we observed not only these glycans, but also highly sialylated and fucosylated structures, and surprisingly glycopeptides with core 4 O-glycans (**Figure S16**). This suggests that when searching MS data, larger glycan databases might be necessary to encompass all of the glycan structures displayed on the complex family of mucin proteins.

### MD simulations of TIM-3 and −4 elucidate the structural and dynamical impact of glycosylation

Following the initial glycoproteomic mapping, we asked how these extremely different glycosylation patterns could affect the structures, and potentially functions, of the TIM proteins. While the IgV domain structures have been solved via X-ray crystallography and NMR, the mucin domains were excluded from analysis, presumably due to the high heterogeneity and density of O-glycosylation.^68,69^ After failed attempts to perform cryoEM on the full TIM-3 and −4 ectodomains, we reasoned that molecular modeling and MD simulations could be an alternative method to predict mucin domain structure and better understand how glycosylation contributes to dynamic properties of these proteins. As described above, the microheterogeneity of each glycosite was incredibly high; thus, in order to accurately reconstruct the TIM glycoproteomic landscape, we needed to identify the most abundant glycan at each residue. To do so, we generated extracted ion chromatograms (XICs) of every glycopeptide detected from TIM-1, −3, and −4 to calculate area-under-the-curve relative quantitation using Thermo Xcalibur (**SI Tables 4, 7-8**). The relative abundance of every detected glycan, at each O-glycosite, is depicted in **Figure 4A** (right). The density and heterogeneity of glycosylation are apparent in TIM-1 and −4, but less so in TIM-3. Additionally, as others before have suggested, the glycan size and heterogeneity was much lower in areas of dense glycosylation; sparse O-glycosites afforded larger and more diverse glycan structures.^70,71^ By obtaining the most abundant O-glycan at each glycosite, we built two all-atom computational models of the fully glycosylated transmembrane glycoproteins TIM-3 and TIM-4 (**Figure 4A**, left) to better understand the contribution of glycan density on the overall flexibility and length of TIMs. Each of these systems contained their respective globular IgV domain, mucin-domain, alpha helical transmembrane domain, and cytoplasmic tail. Approximately 700 and 830 ns of simulation data were generated for TIM-3 and −4, respectively (see SI Methods for full simulation details).

**Figure 4.**
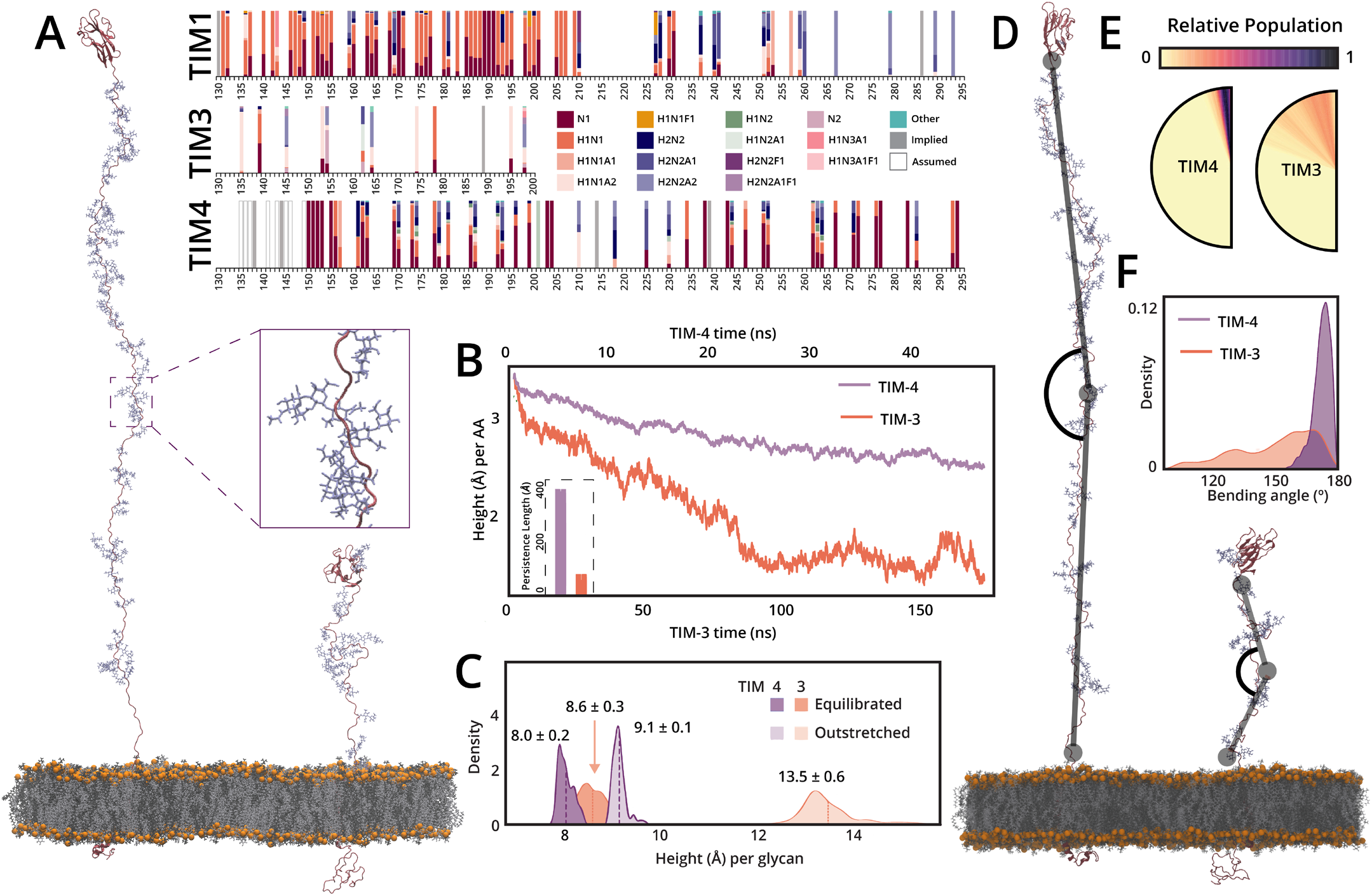
MD simulations of TIM-3 and −4 elucidate the structural and dynamical impact of glycosylation. (A, right) XICs were generated for each glycopeptide from TIM-1, −3, and −4 and area-under-the-curve quantitation was performed; glycan composition color legend shown in center. N - HexNAc (GalNAc), H - hexose (galactose), A - NeuAc (sialic acid), F - fucose. (A, left) Image detailing TIM-4 and TIM-3 models along with an inset view highlighting the dense TIM-4 glycosylation. (B) End-to-end distance of TIM-3 and TIM-4 mucin domains normalized by total length (number of amino acids, AA) within the mucin domains, plotted as a function of simulation length. (inset) Persistence length calculated for TIM-3 and TIM-4 from all simulation replicas. (C) Histograms detailing the end-to-end distance of TIM-3 and TIM-4 mucin domains, normalized by total number of glycans, in the outstretched (starting) conformation (lighter distributions) and equilibrated conformation (darker distributions). (D) Image demonstrating the “bending angle” as calculated in the following panels. (E) Semi-circles graphically detailing the bending angles visited over the complete course of simulations for TIM-3 and −4, angles colored according to relative population. (F) Histograms detailing bending angles sampled by TIM-3 and −4 over the course of all simulations.

To identify the degree to which TIM-3 and TIM-4 mucin domains compress during simulation, we calculated a normalized end-to-end distance of each protein’s mucin domain as a function of time (**Figure 4B**). From these results, we see that TIM-3’s mucin domain, with only 14 glycans and 71 amino acids, compresses far more significantly than TIM-4’s mucin domain, containing 51 glycans and 179 amino acids. We quantified this change by calculating persistence length, which was defined as the distance, in angstroms, at which the motions of two monomers along a polymeric chain become decorrelated from one another. Strong intramolecular interactions within monomers can lead to highly correlated motions along the polymeric chain, overcoming energetic gains of interactions with solvent or enhanced conformational degrees of freedom, and thus long persistence lengths. Using data from our MD simulations, we calculated the persistence lengths of the TIM-3 and −4 mucin domains to be 81 ± 24 Å and 415 ± 10 Å, respectively. Thus, these two mucin-domains have drastically different degrees of correlation within their protein backbones, likely originating from their varied degrees of glycosylation (**Figure 4B,** inset).

During initial analysis and trajectory visualization, we noticed the TIM-3 mucin domain underwent a significant degree of bending such that the IgV domain tilted toward the membrane. To better quantify this, we calculated the angle between two vectors for both TIM-3 and −4 mucin domains: one drawn from the central residue up to the first residue, and another drawn from the central residue down to the last residue (**Figure 4D,** see SI Methods for complete details). We observed that the TIM-4 mucin domain largely sampled bending angles close to 180°, i.e., the TIM-4 mucin domain was largely linear and bottle-brush like. The TIM-3 mucin domain, however, bent quite significantly and sampled a large range of angles with similar probabilities (**Figure 4E-F**).

The mucin domains of TIM-3 and −4 are variably dense in terms of glycosylation. To investigate this effect on mucin identity and functional dynamics, we aimed to quantify total versus effective glycosylation in a thoroughly glycosylated mucin domain (as in TIM-4) versus in a sparsely glycosylated mucin domain (as in TIM-3). Herein, we define the total glycosylation rate as the ratio of the length of the outstretched, unequilibrated mucin domain protein backbone to the total number of glycans. Similarly, we define effective glycosylation as the ratio of the length on a relaxed, equilibrated mucin domain protein backbone to the total number of glycans. These two values thus illustrate the “height per glycan (Å)” under outstretched and relaxed conditions. As shown in **Figure 4C**, the height per glycan in heavily glycosylated TIM-4 remains nearly the same in both outstretched and equilibrated states: 9.1 ± 0.1 Å and 8.0 ± 0.2 Å, respectively. This indicates that upon relaxation of the mucin domain protein backbone, O-glycans still maintain a similar distribution relative to one another as in the fully outstretched case, i.e., total glycosylation equals effective glycosylation. However, for TIM-3, the height per glycan distance drops significantly following equilibration, going from 13.5 ± 0.6 Å to 8.6 ± 0.3 Å. In fact, following equilibration, this height per glycan distance seen in TIM-3 becomes similar to those distributions seen in TIM-4. Through trajectory visualization, specific glycan-glycan pairs in TIM-3 were found to be responsible for a large portion of the decrease in height per glycan distance. Specific distant pairs (≥3 glycans away from one another) of glycans are seen to interact via hydrogen bonding, almost as if these glycans are “holding hands,” as exemplified by the TIM-3 glycan pair G7 and G4 (glycosylation sites T145 and T162, respectively; **Figure S17**). These results demonstrate the power of MD simulations in characterizing members of the mucinome, as glycoproteomic mapping alone cannot provide atomic-level structural insight, including conformational changes that may allow distant O-glycan pairs to find each other and reach new, functionally significant conformations. These combined methods thus have the potential to classify proteins more rigorously within the mucinome.

## Discussion

Historically, numerous challenges have impeded the study of mucin-domain glycoproteins; however, new tools continue to be introduced to unveil mucin glycosylation status, functional roles, and biological impact. Here, we present an addition to this toolkit and use it to better understand mucin-domain glycoprotein structure and dynamics. We first thoroughly characterized the mucinase SmE, which demonstrated a uniquely broad cleavage motif and outperformed commercially available O-glycoproteases, thus enabling unprecedented glycoproteomic mapping of biologically relevant mucin proteins. In particular, we elucidated the glycosylation landscape of clinically relevant immune checkpoint proteins TIM-1, −3, and −4 and used this information to enable glycoproteomic-guided MD simulations for the first time. The data afforded by SmE treatment, in concert with MD simulations, has opened the door to a new realm of atomic-level insight into mucin-domain containing proteins, their structure-function relationships, and their recognition mechanisms by bacterial mucinases. Interplay between glycoproteomic data and molecular modeling offers the potential to expand upon benchmarks for determining mucinome membership, such as mucin-specific persistence length, effective glycosylation, and flexibility. Ultimately, we developed a powerful workflow to understand detailed molecular structure and guide functional assays for all members of the mucinome.

That said, we have only begun to unlock the potential of this workflow, especially as pertained to the TIM family of proteins. To be sure, aberrant glycosylation is a hallmark of cancer and typical O-glycosylation changes involve truncation of normally elaborated glycan structures.^72^ These shortened glycans could strongly impact the “linearity” of the mucin backbone, thus changing TIM protein protrusion from the glycocalyx. As such, transformed O-glycosylation could have implications in how the glycoproteins interact with each other, their ligands, and as a result, intracellular signaling and T cell cytotoxicity. Relatedly, this could greatly influence the efficacy of the 10 anti-TIM-3 antibodies currently being investigated in at least 26 clinical trials (clinicaltrials.gov). Future efforts will be devoted to glycoproteomic mapping of endogenous TIM proteins from primary T cells and patient samples to discover how glycan structures change in health and disease. In concert with biological assays, we will use this information to drive MD simulations that probe how altered glycosylation could affect ligand binding, intracellular interactions, and antibody recognition. Beyond the TIM family of glycoproteins, many other glyco-immune checkpoints have emerged as prominent mechanisms of immune evasion and therapeutic resistance in cancer;^33^ we envision that our workflow will also help elucidate structure-function relationships in these proteins.

Aside from MS analysis, SmE encompasses the potential to make a larger impact on the field of glycobiology. In our previous work, we used StcE for clearance of mucins from the cell surface and determined that Siglec-7, but not Siglec-9, selectively bound to mucin-associated sialoglycans.^19^ We then upcycled an inactive point mutant of StcE to develop staining reagents for Western blot, immunohistochemistry, and flow cytometry.^20^ We also took advantage of the mutant StcE to develop an enrichment procedure that allowed for the selective pulldown of mucin glycoproteins.^10^ Finally, and most recently, an engineered version of StcE conjugated to nanobodies was used for targeted degradation of cancer-associated mucins.^56^ Given that SmE is similarly active on live cells, has a complementary cleavage motif, two mucin binding domains, and potentially different endogenous targets, future work will be aimed at investigating whether SmE can augment our current mucinase toolkit and therapeutic strategies.

Previously, we developed a “Mucin Domain Candidacy Algorithm” to help identify proteins that have a high probability of bearing a mucin domain.^10^ While we recognized at the time that our definition of a mucin domain was novel but rudimentary, the present work has confirmed that our understanding of mucin domains is incomplete. TIM-1 and −4 were predicted by our algorithm to be high confidence mucins, whereas TIM-3 received a score of 0. Here, our glycoproteomic mapping combined with MD simulations demonstrated that absolute glycosylation (i.e., O-glycans per amino acid residue) can be dramatically different than effective glycosylation (i.e., O-glycosylation in relation to total surface area after folding). Thus, while density of O-glycosylation can absolutely be an indication that a mucin domain is present, it does not reveal the entire story, and our definition of a mucin domain continues to develop. While it would be ideal to obtain high-resolution structures of these mucin-domain glycoproteins, that is likely a long-term objective. In the meantime, our workflow is a tangible mechanism for visualizing the enigmatic mucin family to not only better understand the definition of a mucin domain, but to also study the structural dynamics that lie within.

Our ultimate objective is to unravel the complex molecular mechanisms behind glycan structures, patterns, and overall biological functions of mucin domains, but much remains to be accomplished. That said, this work serves as a significant advance toward that overall goal and will find use in furthering our understanding of the mucinome.

## Acknowledgements

We thank Jeffrey Shabanowitz for his technical expertise and thoughtful conversations during the preparation of this manuscript. We also would like to acknowledge the Keck Biophysical Resource for their assistance in cell viability assays. J.C. is supported by a University Fellowship through Yale University; A.S. is supported by the National Institutes of Health Chemical Biology Training Grant (T32 GM067543); K.E.M. and T.M.L. are supported by Yale Endowed Postdoctoral Fellowships in the Biological Sciences. M.A.R. and F.L.K acknowledge the San Diego Supercomputer at UCSD for providing HPC resources that have contributed to the research results reported within this paper. R.E.A. acknowledges support from NIH GM132826, NSF RAPID MCB-2032054, a UC San Diego Moores Cancer Center 2020 SARS-COV-2 seed grant, and U19-AI171954 from NIAID. M.A.R. is supported by NIH T32 EB009380. This work was also supported by a Sarafan ChEM-H Physician-Scientist fellowship (to M.A.H.), the Stanford Maternal & Child Health Research Institute Instructor K Award Support Program (to M.A.H.), a National Blood Foundation Early Career Scientific Research Grant (to M.A.H.), a NIH NHBLI Pathway to Independence award (1K99HL156029-01 to M.A.H.), and NIH grant R01CA200423 (to C.R.B.). D.J.S. was supported by a US NSF Graduate Research Fellowship and Stanford Graduate Fellowship M.J.F. was supported by a University of Redlands Faculty Research Grant. SAM is supported by the Yale Science Development Fund and a NIGMS R35-GM147039.

## Author Contributions

S.A.M., R.E.A., and C.R.B. advised the project and oversaw experiments. S.A.M. and K.E.M. designed mass spectrometry experiments. J.C., A.D.S., K.E.M., T.M.L., and D.I. performed mass spectrometry and cell biology experiments. J.C., A.D.S., K.E.M., D.I., A.S.B., V.A., and C.K. analyzed mass spectrometry data. M.J.F. performed molecular docking experiments and analyzed associated data. M.A.H, D.J.S., and K.H.T. provided materials and plasmids. M.A.R., F.L.K., and R.E.A. designed all MD simulations. M.A.R. constructed TIM-3, TIM-4 models and conducted MD simulations. M.A.R. and F.L.K. conducted analysis from MD simulations. S.A.M., J.C., A.D.S., T.M.L., K.E.M., M.J.F., M.A.R., and F.L.K. wrote the manuscript with additional input from all authors.

