## Supplemental Information for "Glycoproteomic landscape and structural dynamics of TIM family immune checkpoints enabled by mucinase SmE"

**This PDF file includes:**

Supplementary Figures 1 to 18
Supplementary Materials and Methods
Supplementary References

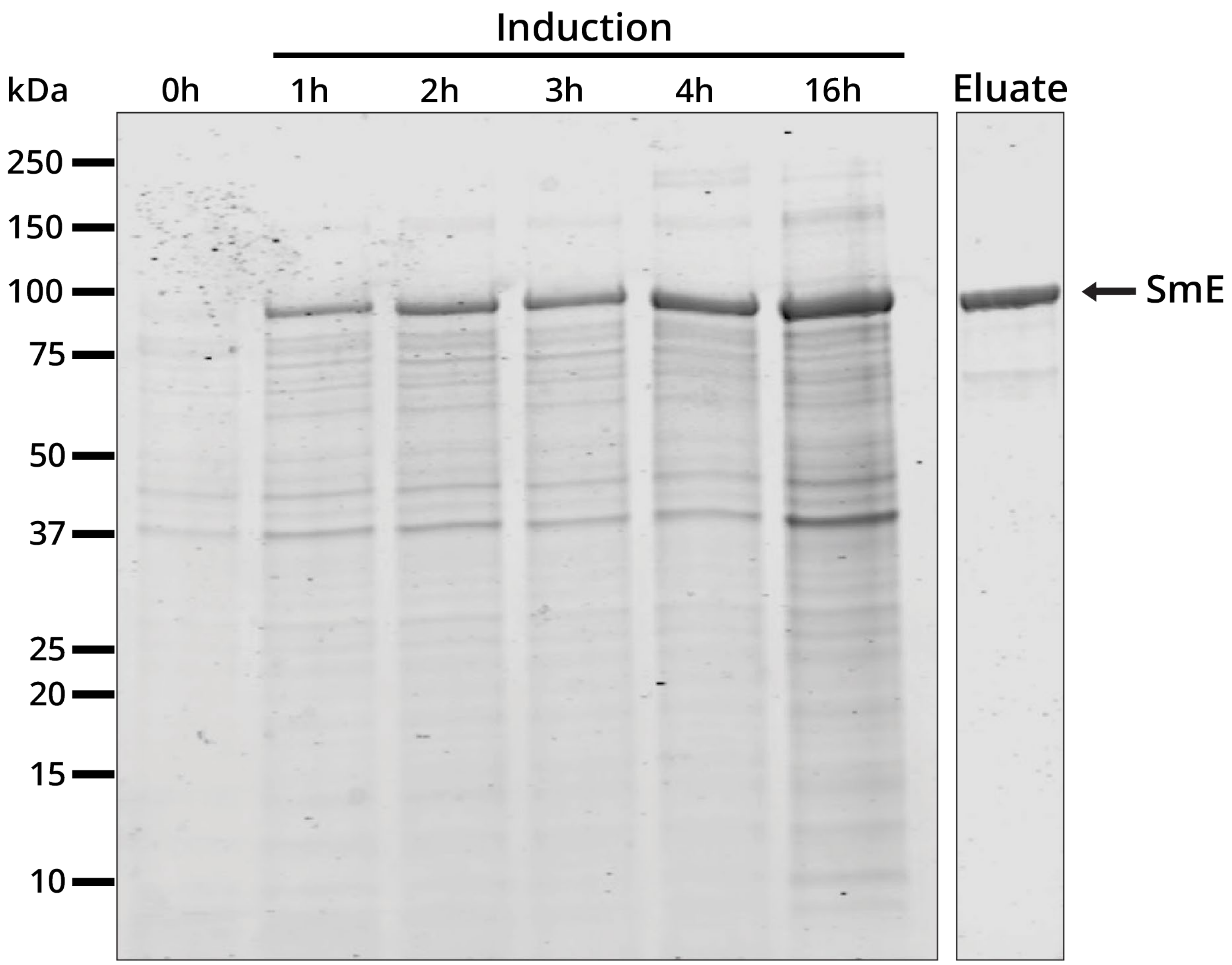


**Figure S1. Expression and purification of SmE.** SmE was expressed overnight at 16 °C. SDS-PAGE gel was stained with Coomassie (Bulldog-Bio) and imaged using an Odyssey CLx Near-Infrared Fluorescence Imaging System (LI-COR Biosciences). SmE ran at the predicted molecular weight of 94 kDa, as indicated by the arrow.

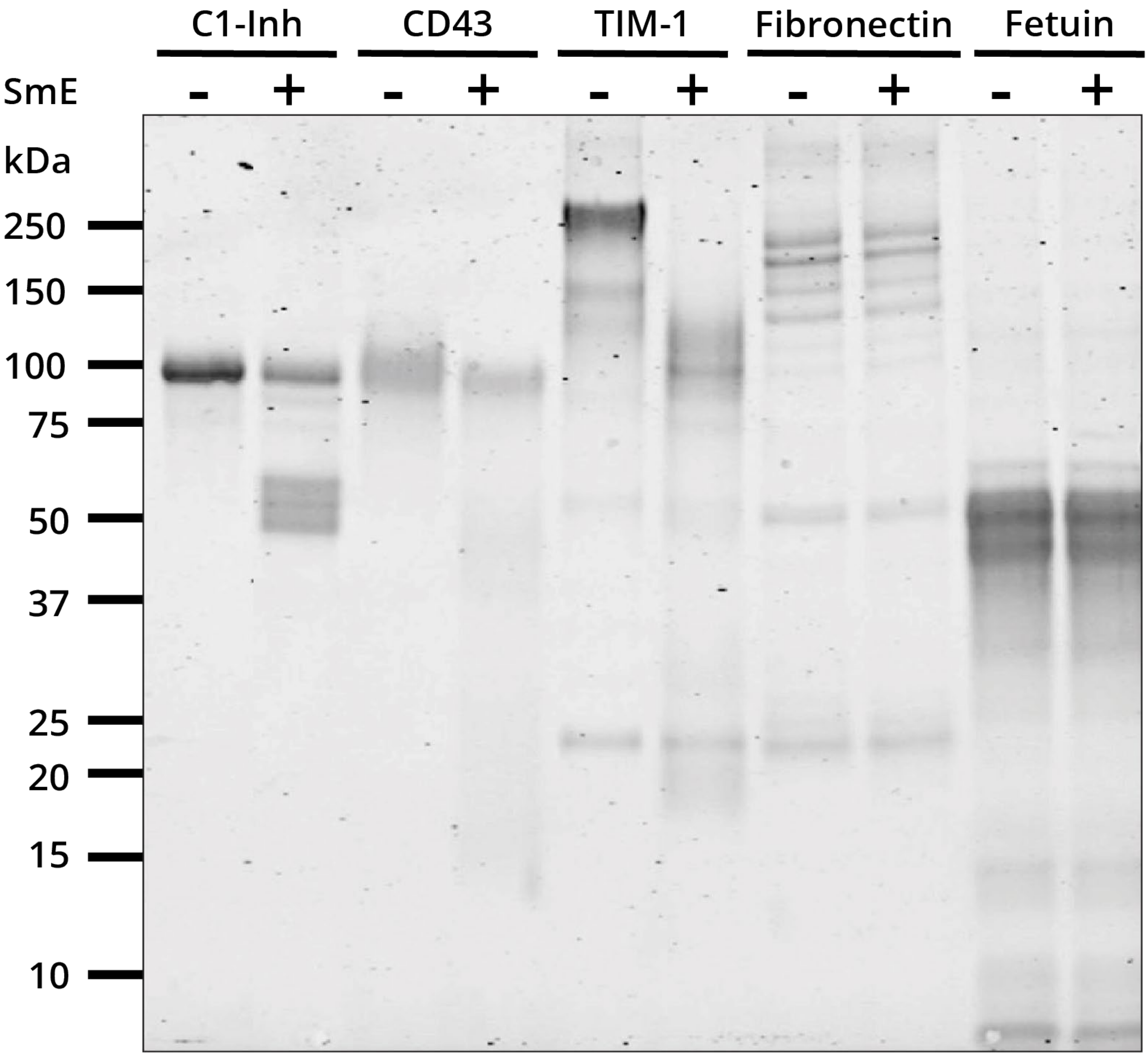


**Figure S2. SmE selectivity for mucin-domain glycoproteins.** The recombinant proteins shown were incubated with SmE at a 1:20 E:S ratio overnight at 37 °C and the digests were separated by SDS-PAGE. SDS-PAGE gel was stained with Coomassie (Bulldog-Bio) and visualized on an Odyssey CLx Near-Infrared Fluorescence Imaging System (LI-COR Biosciences).

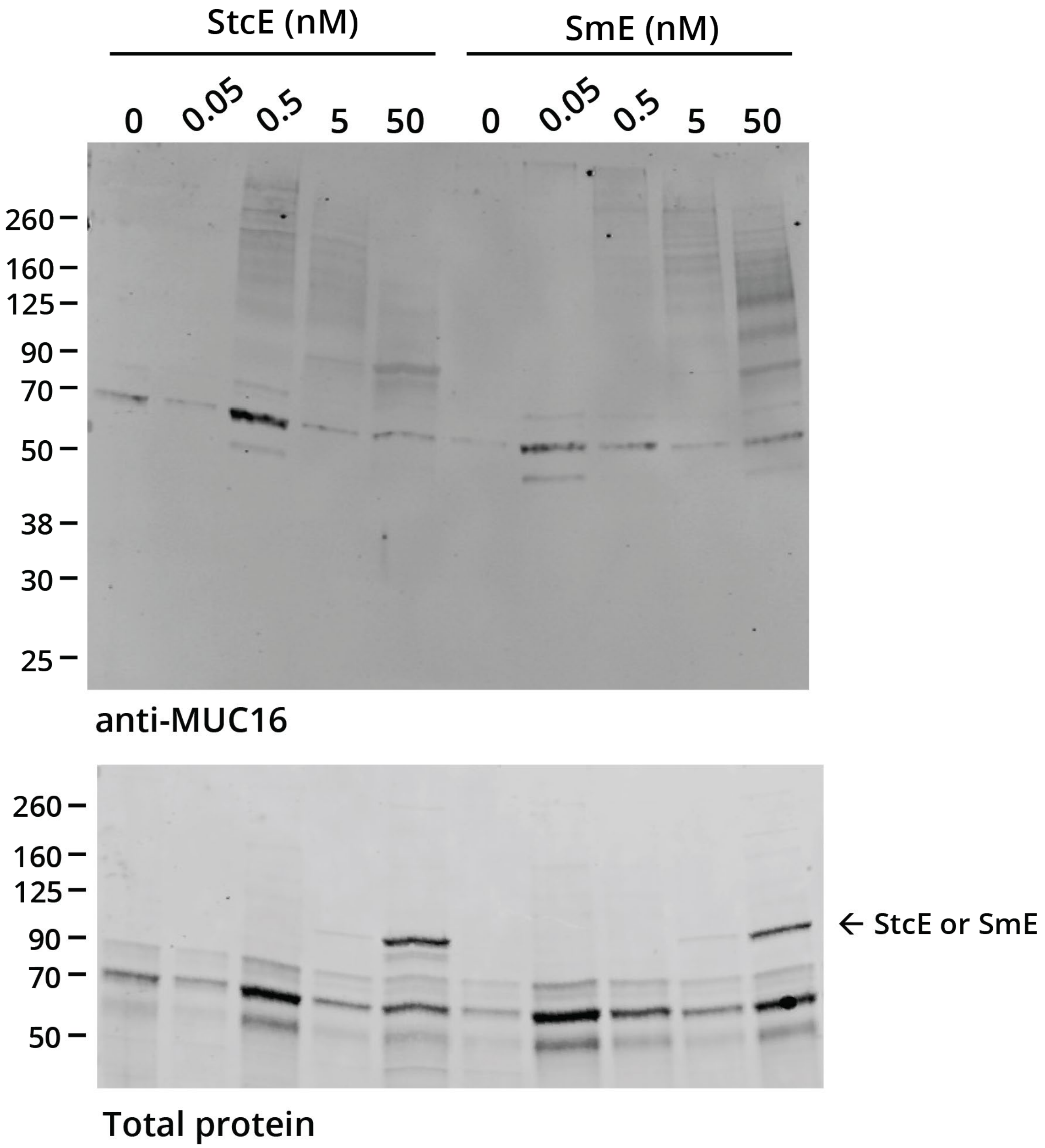


**Figure S3. MUC16 is released into cell supernatant after mucinase treatment**. HeLa cells were treated with StcE (left) or SmE (right) at the noted concentrations for 60 min. Following treatment, the supernatant was transferred to 15 mL conical tubes containing 75 µL of EDTA to stop enzyme processing. The supernatants were then concentrated using 3 kDa MWCO spin filters and then diluted with 4X LDS sample buffer to a final concentration of 1X. The samples were then boiled at 95 °C for 5 min and loaded onto a 4-12% Bis Tris gel for separation by gel electrophoresis and probing for MUC16 via Western blot. Proteins were transferred to a nitrocellulose membrane using the Trans-Blot Turbo Transfer System (Bio-Rad) at a constant 2.5 A for 15 min. Total protein was quantified using REVERT stain before primary antibody incubation overnight at 4 °C. An IR800 dye labeled secondary antibody was used according to manufacturer's instructions for visualization on a Licor Odyssey instrument.

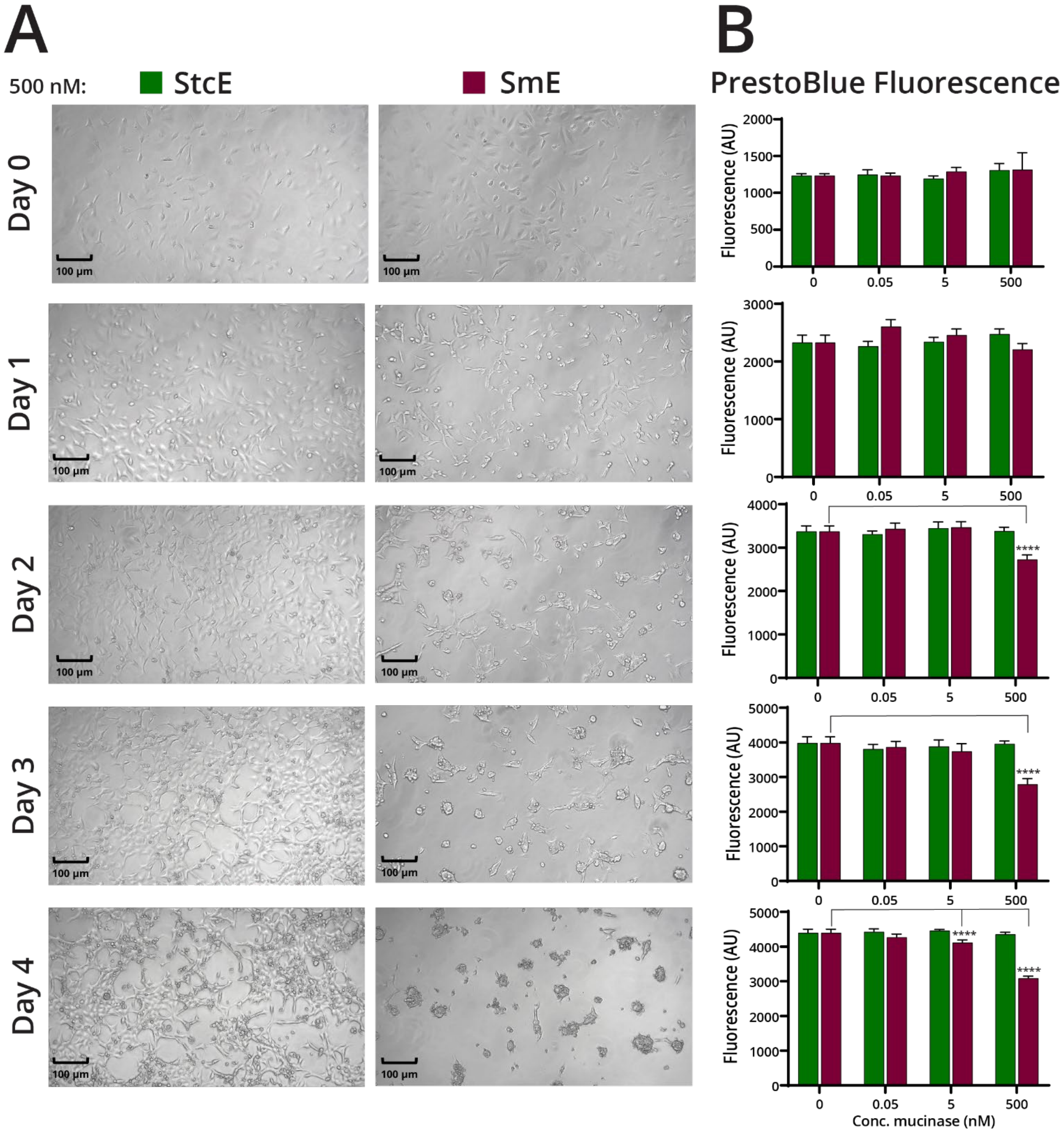


**Figure S4. SmE treatment was nontoxic to HeLa cells at moderate concentrations and/or durations.** (A) Live cell microscopic images of HeLa cells treated with 500 nM StcE or SmE, taken every 24 hours for a total of 4 days. (B) Cellular viability was measured using a resorufin-based dye (PrestoBlue, Thermo Fisher Scientific), at 0, 0.05, 5 and 500 nM StcE or SmE treatment over 4 days. Green: StcE, maroon: SmE. Statistical significance was determined using the two-way ANOVA analysis in Graphpad PRISM software and is reported with respect to the no mucinase control condition. **** indicates a p value <0.0001. Scale bar = 100 µm.

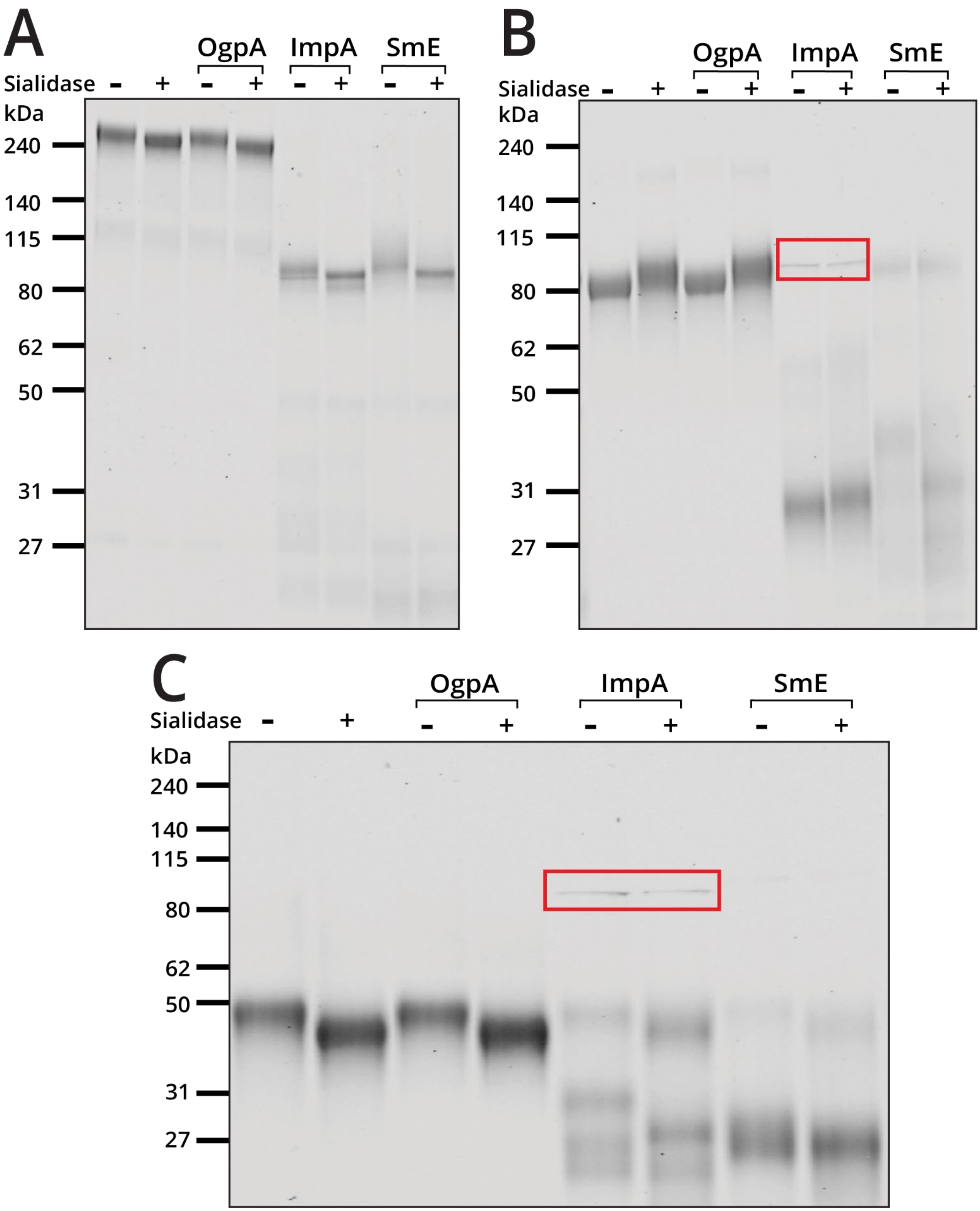


**Figure S5**. **Benchmarking OgpA, ImpA, and SmE activity on TIM proteins.** Recombinant proteins (A) TIM-1, (B) TIM-4 and (C) TIM-3 were reacted with O-glycoproteases with and without sialidase overnight at 37 °C. For SmE digestion, we employed a 1:10 E:S ratio, while for OgpA and ImpA, manufacturer's recommended conditions were used. All digests were separated via SDS-PAGE and stained with Coomassie (Bulldog-Bio). Gels were imaged on an Odyssey CLx Near-Infrared Fluorescence Imaging System (LI-COR Biosciences). ImpA is denoted by the red box.

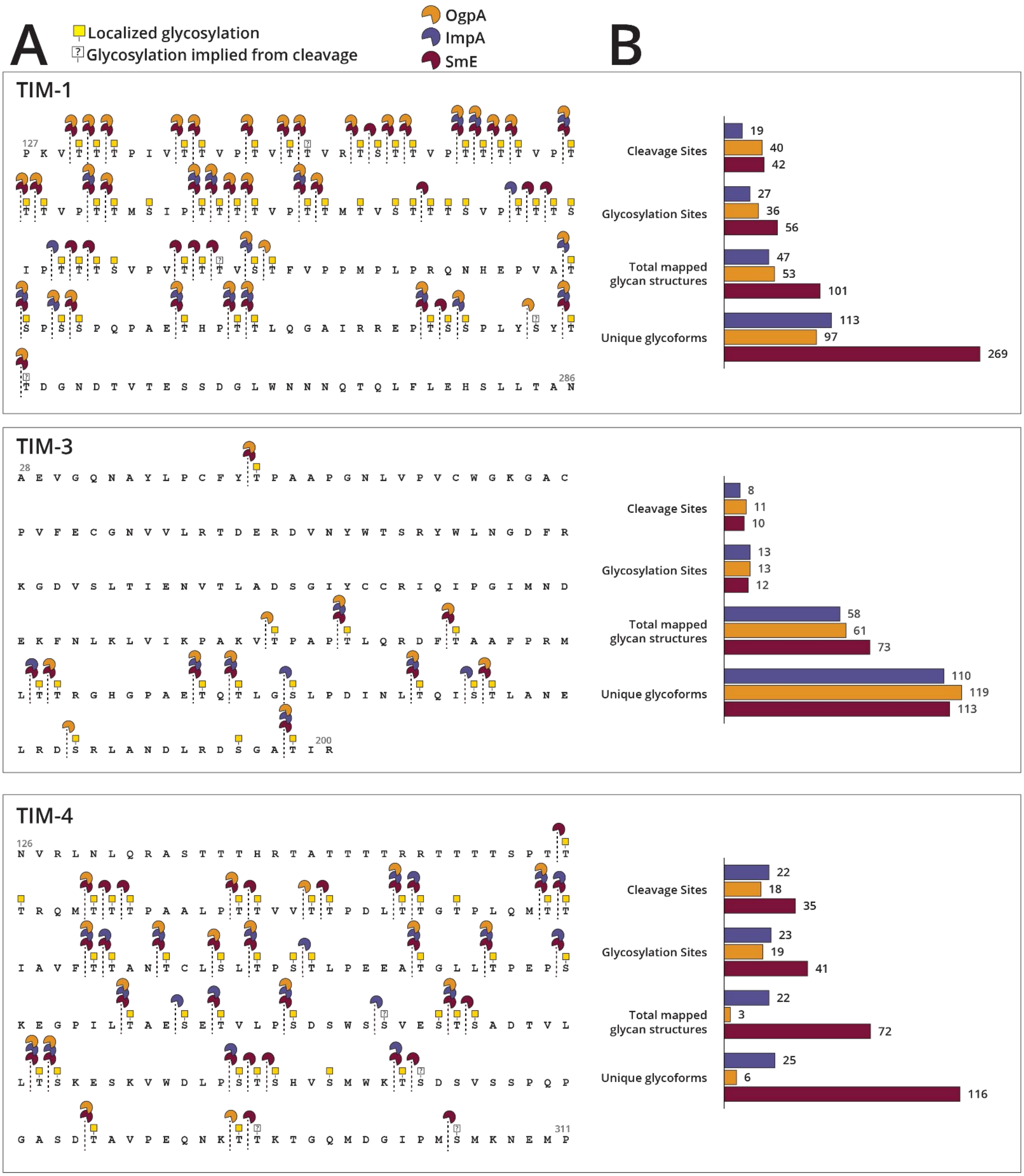
A
Localized glycosylation
Glycosylation implied from cleavage
OgpA
ImpA
SmE
B
TIM-1
Cleavage Sites
19
40
42
Glycosylation Sites
27
36
56
Total mapped glycan structures
47
53
101
Unique glycoforms
113
97
269
TIM-3
Cleavage Sites
8
11
10
Glycosylation Sites
13
13
12
Total mapped glycan structures
58
61
73
Unique glycoforms
110
119
113
TIM-4
Cleavage Sites
22
18
35
Glycosylation Sites
23
19
41
Total mapped glycan structures
22
3
72
Unique glycoforms
25
6
116

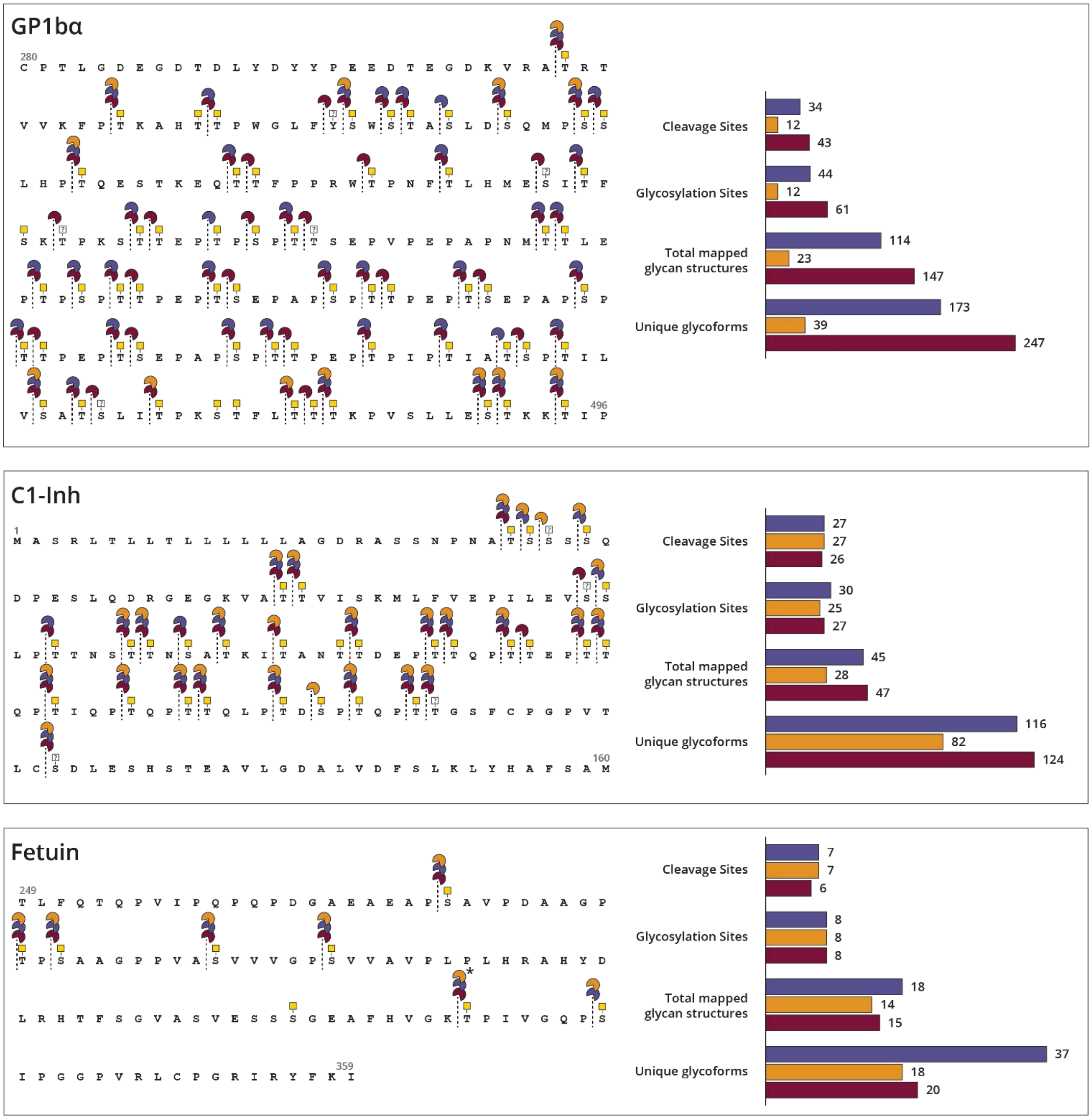


**Figure S6. Total cleavage events.** (A) Cleavage maps depict observed glycosites and cleavage sites of each enzyme from MS analysis. (B) Graphical interpretation of individual glycoprotein cleavage sites, glycosites, and glycoforms identified by treatment with each enzyme. Red: SmE, blue: ImpA, yellow: OgpA. Asterisk in fetuin cleavage map indicates cleavage attributable to either mucinase or co-enzyme (trypsin); white squares with question marks indicate implied glycosites with observed cleavage.

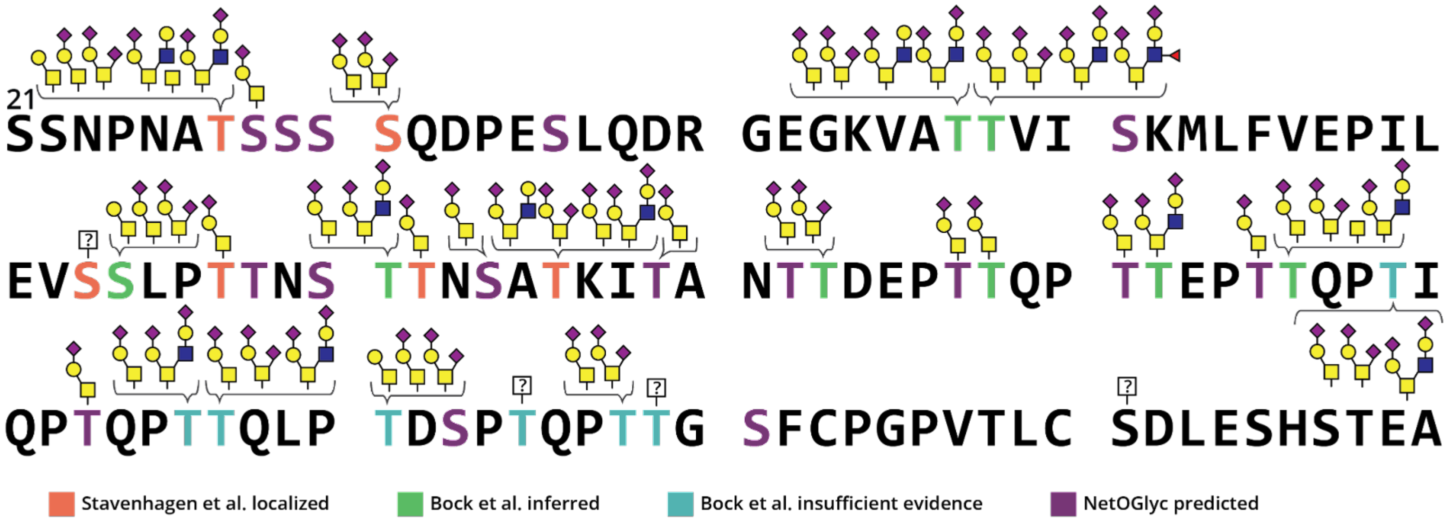


**Figure S7. C1-Inh glycoproteomic landscape.** C1-Inh isolated from human plasma was subjected to digestion with SmE and trypsin followed by MS analysis and manual data validation. Depicted are residues 21-140 which comprise the C1-Inh mucin domain. Glycans depicted in brackets were detected on the associated residues. The colored residues were either detected (orange; Stavenhagen *et al.*),[1] inferred (green; Bock *et al.*),[2] hinted at without sufficient evidence (blue; Bock *et al.*),[2] or predicted by NetOGlyc 4.0 (purple).

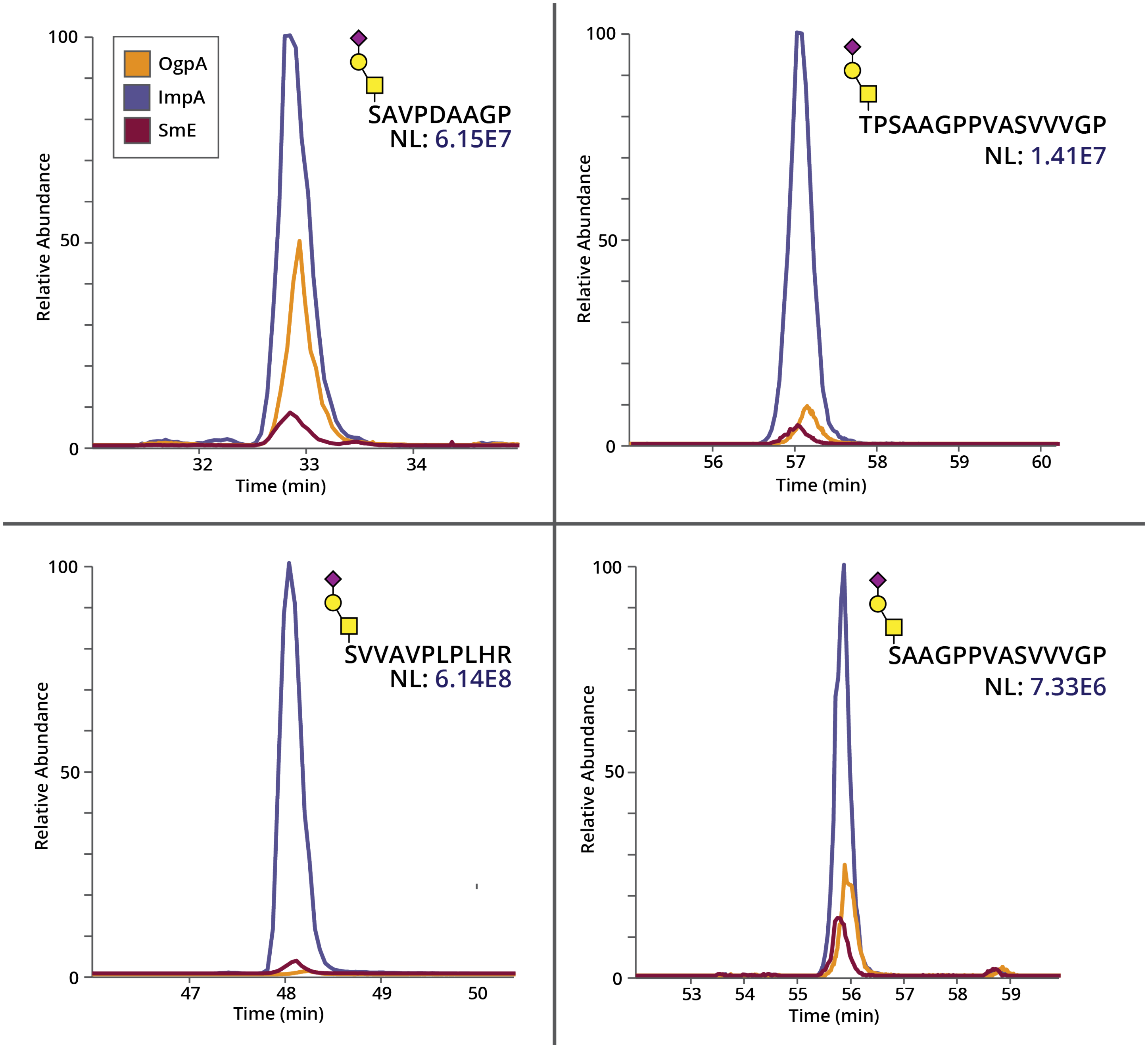


**Figure S8. Extracted ion chromatograms of cleaved glycopeptides from fetuin**. Using Thermo Xcalibur, extracted ion chromatograms (XICs) were generated for four glycopeptides from the OgpA, ImpA, and SmE digest of fetuin. The XICs of specific glycopeptides were normalized to indicate relative abundance in each analysis. Yellow: OgpA, blue: ImpA, red: SmE.

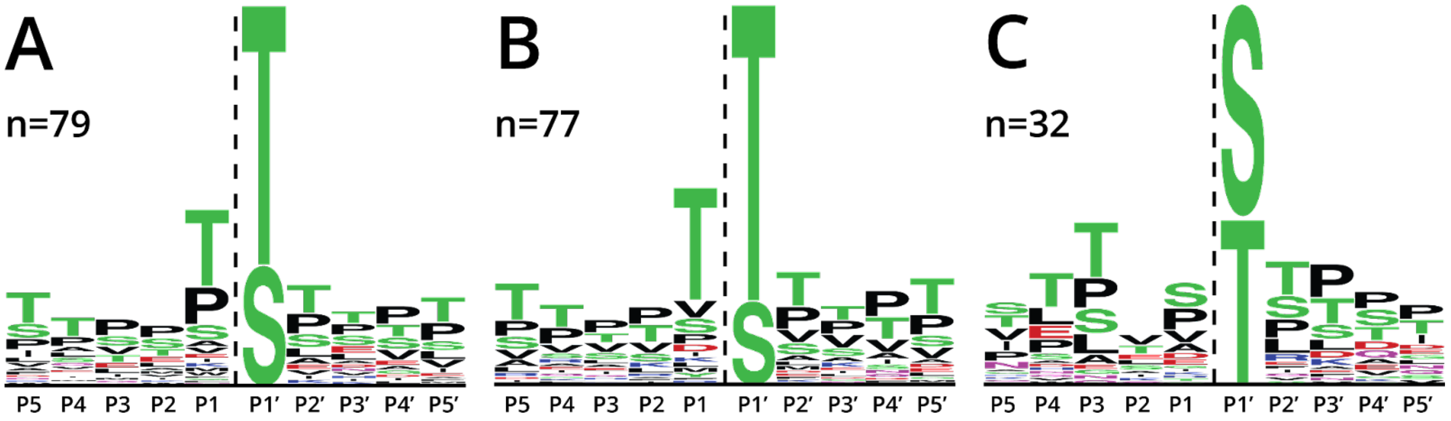


**Figure S9. Anti-logos demonstrate missed cleavage events and less preferred P1/P1' residues.** Using cleavage maps from Figure S5, all sites without observed cleavage were loaded into weblogo.berkeley.edu. OgpA (A) and ImpA (B) demonstrated less efficiency with Thr in the P1 position. SmE (C) uniquely showed lower preference for Ser in the P1' position.

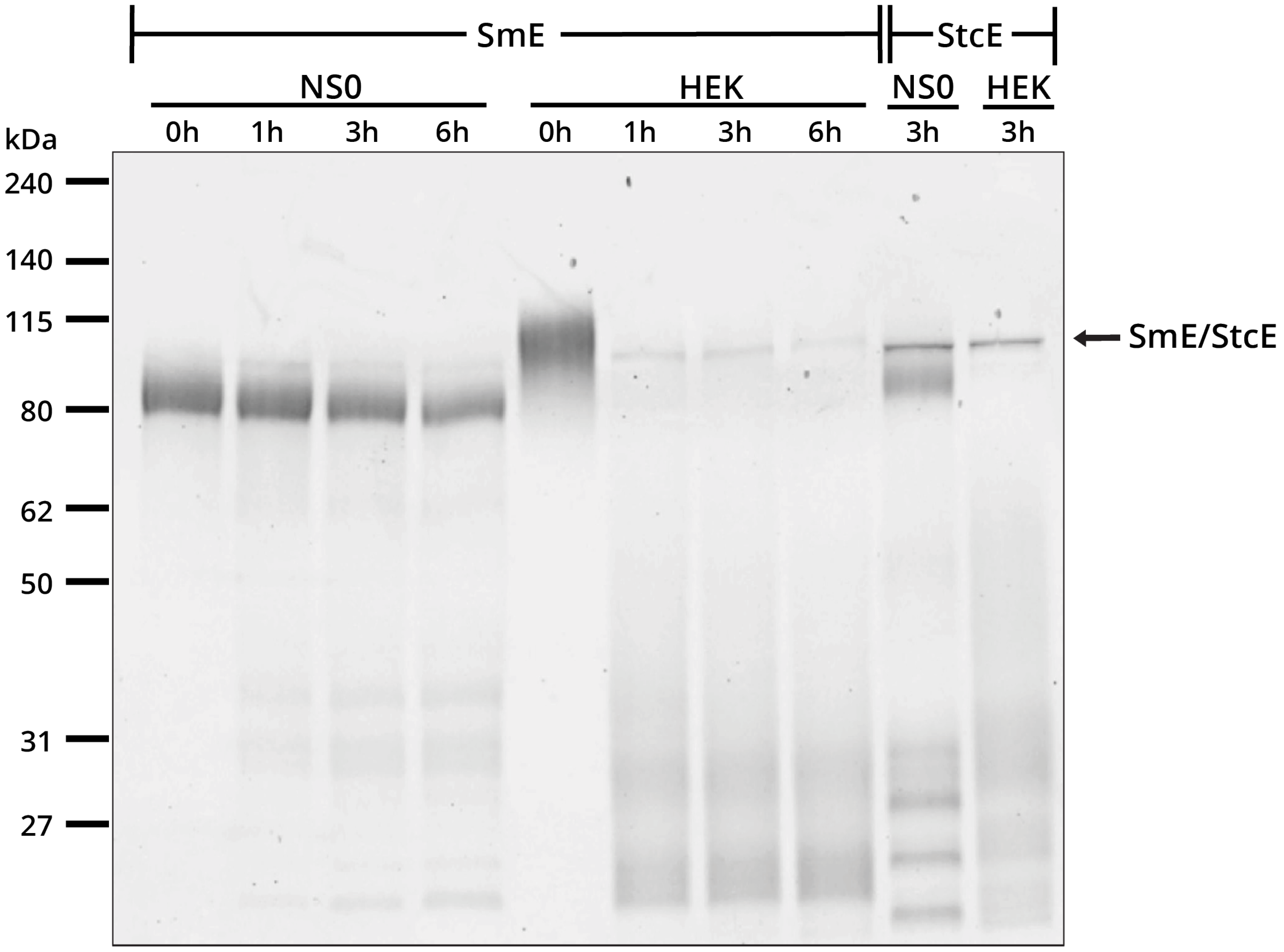


**Figure S10. Comparison of SmE activity on recombinant mouse (NS0) or human (HEK293)-derived TIM-1.** Recombinant human TIM-1 expressed in NS0 or HEK293 cells were reacted with SmE at a 1:10 E:S ratio for 1, 3, or 6 hours at 37 °C and StcE at a 1:10 E:S ratio for 3 hours at 37 °C. All digests were separated by SDS-PAGE and Coomassie stained (Bulldog-Bio). Gel was visualized on an Odyssey CLx Near-Infrared Fluorescence Imaging System (LI-COR Biosciences).

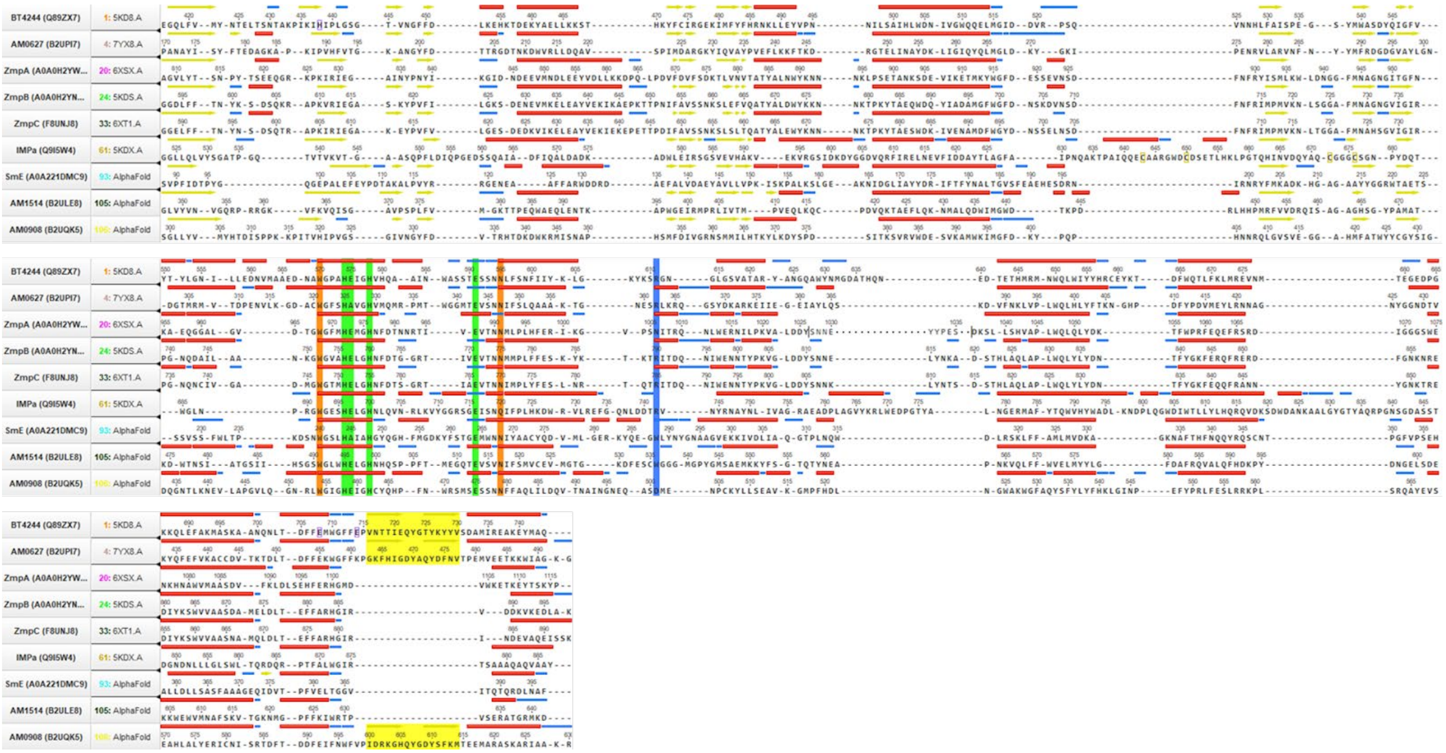

**Figure S11. Sequence alignment of the catalytic PF13402 domains found in characterized O-glycoproteases.** The X-ray crystal structures of BT4244, AM0627, ZmpA, ZmpB, ZmpC, and ImpA as well as the AlphaFold-predicted structures of SmE, AM1514, and AM0908 were structurally overlaid using the conserved zinc-binding and catalytic residues (green). The structural overlays were then used to generate the above sequence alignments, highlighting key secondary structures: alpha helices (red), beta sheets (underlined yellow), and turns (underlined blue). In addition to the catalytic core residues (green), we have highlighted conserved (orange) and semiconserved (highlighted blue) residues involved in recognizing P1’ glycans, as well as residues in a semiconserved beta-hairpin (highlighted yellow) that can potentially recognize P1 glycans. Notably, SmE has neither the semiconserved Arg residue nor the semiconserved beta-hairpin.

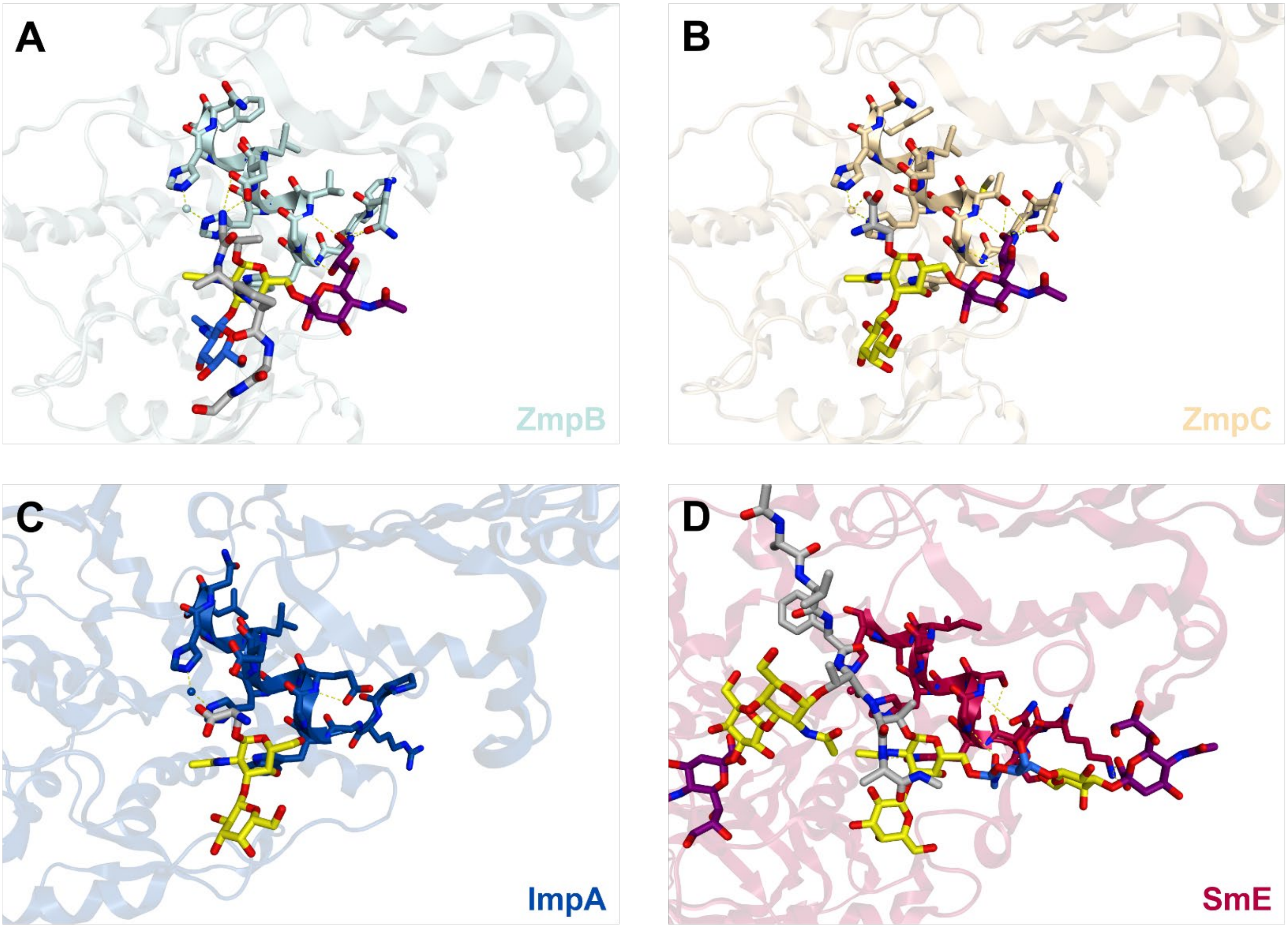


**Figure S12. Ligand-bound crystal structures of O-glycoproteases.** Structures of ligand-bound catalytic helices of (A) ZmpB, (B) ZmpC, and (C) ImpA as well as the modeled structure of (D) SmE. Both ZmpB and ZmpC form specific contacts (yellow dashes) between residues near the terminus of the catalytic helix (colored sticks) and the branching sialic acid residue (purple sticks) found in the ligand at P1'. The ligand in ImpA does not contain sialic acid; analogous interactions can likely form, given the similar length of its catalytic helix, imparting ImpA with the ability to accommodate branched ligands. In the modeled structure, SmE forms contacts with the GalNAc residue (yellow sticks) and additional contacts with the sialic acid residue (purple sticks) of the glycan, which may explain its ability to accommodate larger branched glycans at P1'.

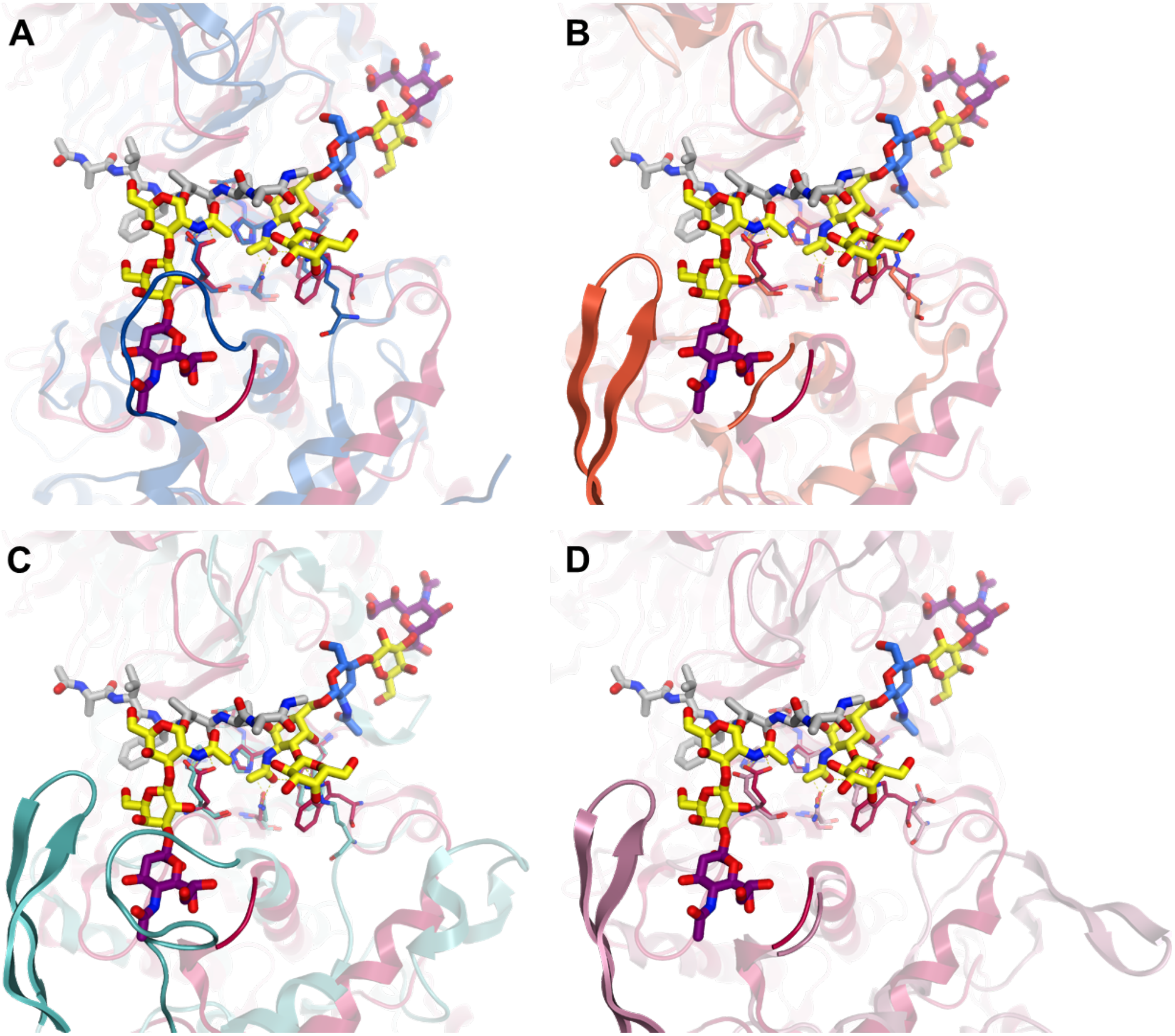


**Figure S13. Structural alignments of O-glycoproteases.** Overlays of the docked SmE-glycopeptide complex (red) with the crystal structures of (A) ImpA (blue), (B) AM0627 (orange), and (C) BT4244 (mint) as well as the AlphaFold predicted structure of (D) AM0908 (pink). The residues (thin sticks) of the catalytic core and the conserved residues that bind the P1' glycan are shown with the docked ligand (thick sticks). The loops and beta hairpins that form the steric environment around the P1 glycan are darkened for emphasis. (A) The loop in SmE is predicted to be short, which could explain why SmE can sterically accommodate and act on substrates with glycosylation at P1; the loop of ImpA is long and likely prevents such activity. (B) AM0627 has a short loop that can accommodate different glycans at P1. (C) The loop of BT4244 is much larger, and this difference could resolve discrepancies reported across the literature and may explain why BT4244 showed preference for engineered substrates bearing the smaller Tn-antigen. (D) AM0908 is predicted to have a short loop similar to AM0627; despite this potential similarity, the two enzymes display different preference for P1 glycosylation, which cannot be explained by this reasoning and may be the result of subtler differences in their sequences.

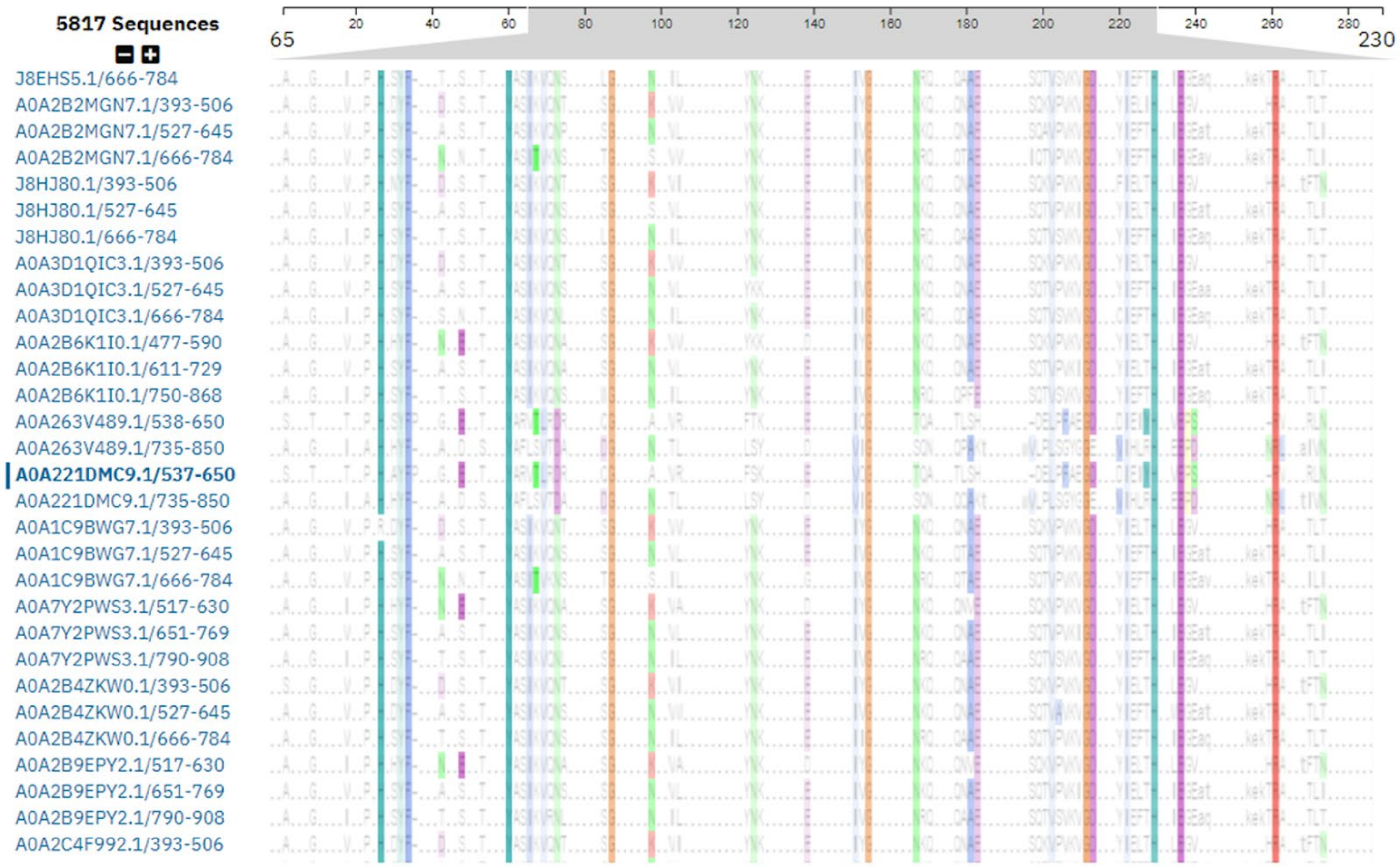


**Figure S14**. **Sequence alignment of PF03272 modules.** Conservation across the 5,817 mucin-binding modules (PF03272) found in proteins, including SmE (A0A221DMC9.1/537-650), listed in UniProt. Degree of conservation is shown using clustal2 coloring, highlighting two conserved motifs (HxxFxxxxY and HxExxR) found in loops clustered together to form a single binding pocket in the AlphaFold structure of SmE. Full results are available at https://www.ebi.ac.uk/interpro/entry/pfam/PF03272/entry_alignments/?type=uniport.

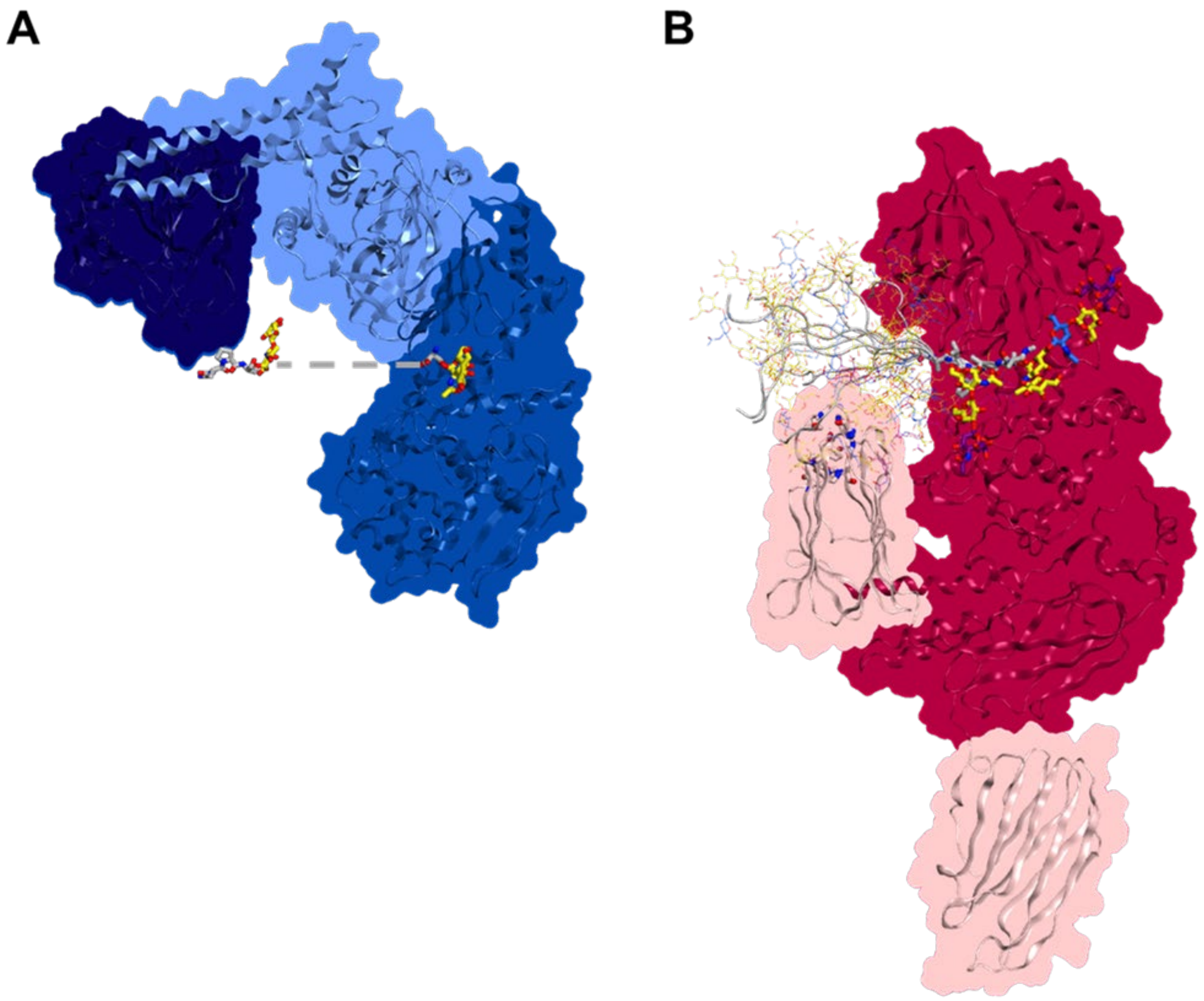


**Figure S15. Potential cooperativity of ligand binding in ImpA and SmE.** (A) Overlay of crystal structures of ImpA with crystallized ligands in the catalytic and accessory domains. The ligand (thick sticks) crystallized in the catalytic domain (blue loops and shaded surface) as well as the ligand (thick sticks) in the accessory N-terminal domain (dark blue loops and shaded surface) are small fragments that likely reflect the orientation of larger mucin-like substrates (dashed line), highlighting potential cooperativity between the different domains in ImpA. (B) Grafted TIM-4 fragments overlaid with the modeled structure of SmE. The catalytic domain (red loops and shaded surface) and the two mucin-binding modules (light pink loops and shaded surface) are shown, including the conserved motifs (HxxFxxxxY and HxExxR, thick sticks) in the loops of the mucin-binding module adjacent to the catalytic site. In addition, the amino acid and glycan residues of the original docked glycopeptide (thick sticks) is displayed, grafted together with ten different TIM-4 fragments. Only the peptide backbone (gray ribbon) and glycans (thin sticks) of these fragments have been highlighted for clarity. Some glycans were found to flank the conserved residues of the mucin-binding module, suggesting that the catalytic and accessory domains of SmE may cooperatively recognize and cleave mucin-like substrates.

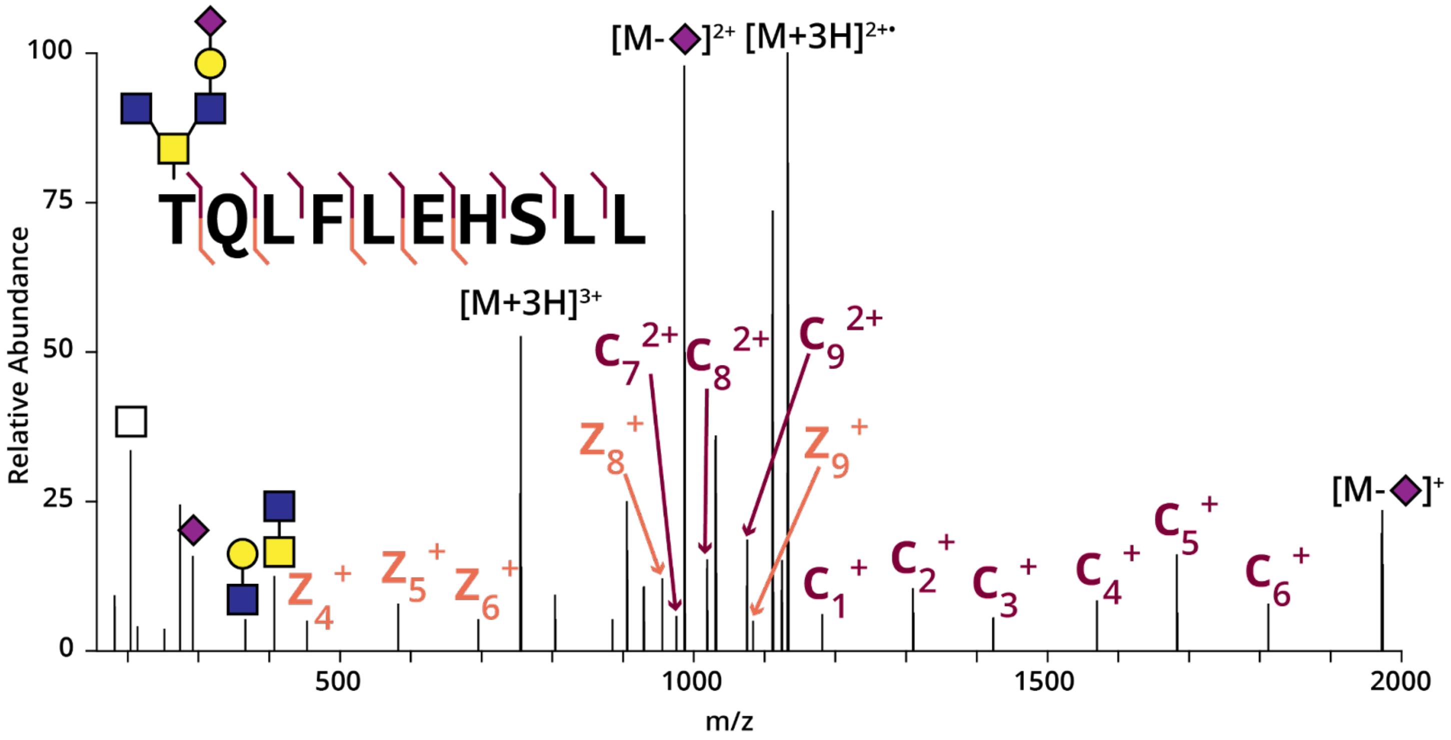


**Figure S16. Recombinantly expressed TIM proteins contain complex glycan structures**. TIM-1 was subjected to digestion with SmE followed by MS analysis and manual data interpretation. An EThcD spectrum from glycopeptide TQLFLEHSLL is depicted above, bearing a core 4 glycan structure. We observed full sequence coverage and unambiguous site-localization.

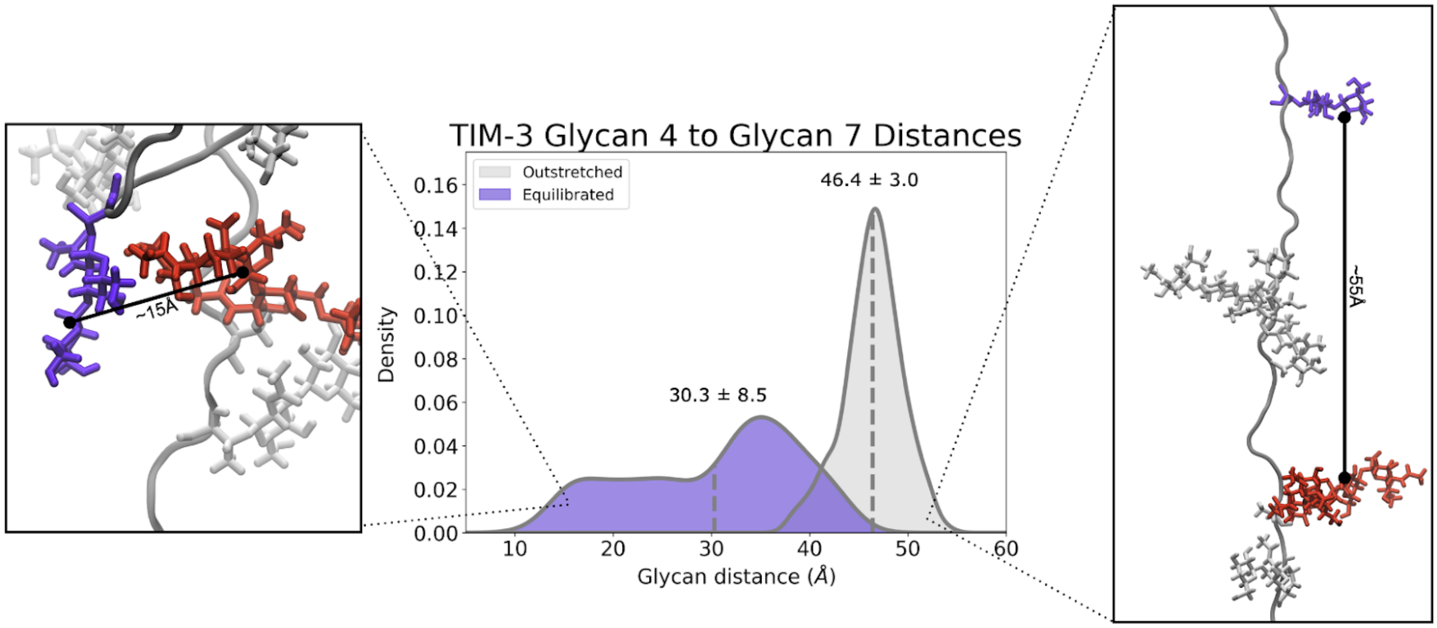


**Figure S17**. **Histogram detailing distance between glycans G4 (site T162) and G7 (site T145) in outstretched versus equilibrated conformation, measured throughout molecular dynamics simulation**. Left panel is a frame taken from an equilibrated conformation with a glycan distance of approximately 15 Å between G4 and G7. Right panel is a frame taken from an outstretched/not fully equilibrated conformation with a glycan distance of approximately 55 Å between G4 and G7.

179 TGTPLQMTTIAVFT|TANTCLSLTPSTLP 206

| Glycan | Occupancy |
|---|---|
| H1N1A1 | 19.6% |
| H1N1 | 18.4% |
| N1 | 15.7% |
| n/a | 12.9% |
| H1N1A2 | 12.0% |
| H2N2 | 9.8% |
| H1N2 | 3.5% |
| H2N2A1 | 3.3% |
| H1N1F1 | 2.9% |
| H1N2A1 | 1.1% |
| H2N2A2 | 0.6% |
| N2 | 0.1% |

| Glycan | Occupancy |
|---|---|
| H2N2A1 | 43.0% |
| N1 | 26.7% |
| H2N2 | 15.3% |
| H1N1 | 8.2% |
| H2N2F1 | 1.9% |
| H1N2 | 1.4% |
| H1N1A2 | 1.4% |
| H1N1A1 | 1.1% |
| H1N2A1 | 0.8% |

**Figure S18. TIM-4 T192 and T193 glycan abundances.** The relative abundance of different glycans branching from T192 and T193 of TIM-4, as revealed through digestion with SmE and subsequent MS analysis. The H1N1A1 and H2N2A1 glycans were the most abundant species at these positions, and were incorporated into the ligand used in docking experiments with SmE.

### Supplementary Materials and Methods

#### Materials

Recombinantly expressed TIM-1 and TIM-4 were purchased from R&D Systems (9319-TM, 9407-TM). For structural characterization, TIM-1 was purchased from R&D systems (11157-TM) and TIM-4 was purchased from LifeSpan Biosciences (LS-G139224). TIM-3 was purchased from LifeSpan Biosciences (LS-G97947). CD43 and TIM-1 recombinantly expressed in NS0 cells were purchased from R&D systems (9680-CD,1750-TM). C1-INH and fibronectin isolated from human plasma were purchased from Sigma Aldrich (E0518, F1056). Bovine Fetuin-A was purchased from Promega (V4961). GP1bα was isolated as described previously.[3] The plasmid for His-tagged pET28a-SmEnhancin and recombinant StcE protein were kindly provided by the Bertozzi laboratory.

#### Expression and purification of SmEnhancin

For pET28a-SmEnhancin plasmid extraction, a 50 µL aliquot of chemically competent *E. coli* DH5α cells (NEB, C2988J) was thawed on ice. Approximately 100 ng of plasmid was added to the cells and incubated on ice for 30 min. Cells were then transformed by heat-shock at 42 °C for 30 seconds. Room temperature SOC media (Invitrogen, 15544-034) was added (950 µL) to the cells then incubated at 37 °C with agitation at 250 rpm for one hour. A 150 µL aliquot of the transformed cells was then transferred to a LB-agar (Fisher, BP1425) plate with kanamycin (Sigma Aldrich, K1377) and incubated overnight at 37 °C. Kanamycin was used throughout at a final concentration of 50 µg/mL. A single colony was picked and used to inoculate an overnight culture of 100 mL Luria broth (LB) (Sigma Aldrich, L3022) with kanamycin. The culture was incubated at 37 °C with agitation at 250 rpm. pET28a-SmEnhancin plasmid DNA was extracted using a Qiagen Plasmid Midi Kit (Qiagen, 12143) using the protocol provided by the manufacturer. DNA concentration was determined by NanoDrop One Microvolume UV-Vis Spectrophotometer (Thermo-Fisher) then stored at -80 °C. After extraction, the DNA sequence was verified using Plasmidsaurus.

For protein expression of SmEnhancin (SmE), a 20 µL aliquot of competent *E. coli* BL21(DE3) cells (Millipore Sigma, 69450-4) was thawed on ice. Approximately 10 ng of pET28a-SmEnhancin was added to cells and incubated on ice for 5 min. Cells were then transformed by heat-shock at 42 °C for 30 seconds. Room temperature SOC media was added (80 µL) to the cells then incubated at 37 °C with agitation at 250 rpm for one hour. A 100 µL aliquot of the transformed cells was then transferred to a LB-agar plate with kanamycin and incubated overnight at 37 °C for colony growth. A single colony was picked to inoculate a 10 mL overnight culture of LB with kanamycin and incubated at 37 °C with agitation at 250 rpm. A glycerol stock was made by mixing 4 mL of the overnight culture with 4 mL of 50% glycerol (Sigma Aldrich, G7893) and stored at -80 °C. The remaining overnight culture was used to inoculate a 1L LB culture with kanamycin. This culture was also incubated at 37 °C with agitation at 250 rpm until it reached an optical density of 0.6-0.8. The maxi culture was then induced with a final concentration of 0.1 mM isopropyl-β-D-1-thiogalactopyranoside (IPTG) (American Bio, AB00841) and grown overnight at 16 °C with agitation at 250 rpm. The bacterial cells were harvested by centrifugation at 3000 *x g* for 45 min at 4 °C. The supernatant was decanted and the cell pellet was stored at -80 °C until lysis was performed.

The cell pellet was lysed in a buffer containing 20 mM Tris-HCl at pH 8 (Thermo Scientific, J3636.K2), 200 mM NaCl (Fisher Scientific, S25877), 2 mM magnesium chloride ($MgCl_2$) (American Bio, AB09006), 10% glycerol, and 125 U/mL Benzonase nuclease (Sigma Aldrich E1014). Additionally, 100 µg/mL lysozyme (Thermo Scientific, 89833) and 1% Triton X-100 (Alfa Aesar, J66624) were added to aid cell lysis. For inhibition of protease activity, a cOmplete Mini EDTA-free protease inhibitor cocktail (Roche, 11836170001) was used alongside 1 mM phenylmethylsulfonyl fluoride (PMSF) (American Bio, AB01620). Cells were then resuspended using 1 mL of chilled lysis buffer per gram of cells. The cell suspension was further homogenized by five pulses of probe sonication with 5 seconds of sonication at 35% amplitude followed by 15 second pauses (QSonica Q500). The solution was kept on ice throughout sonication to prevent protein denaturation/degradation. Lysate was clarified by spinning at 25,000 *x g* for 45 min at 4 °C and the supernatant was filtered sequentially through 0.45 µm (Millipore, SLHAR33SS), and 0.2 µm (Cytiva, 10462300) syringe filters.

The protein was purified using an ÄKTA Pure FPLC (Cytiva) with a HisTrap HP column (Cytiva, 17524801). The column was equilibrated for 5 column volumes (CV) at 1 mL/min using buffer A (20 mM Tris-HCl pH8, 500 mM NaCl, 25 mM imidazole (Sigma Aldrich, I202)) prior to loading the sample at a flow rate of 0.5 mL/min. A conditional wash was then performed, rinsing at 1 mL/min with buffer A until the absorbance of the column flowthrough fell below 10 mAU. The protein was then eluted with a 15 CV linear gradient to 100% solvent B (20 mM Tris-HCl pH8, 500 mM NaCl, 500 mM imidazole). During sample load and wash phases, 5 mL fractions of the eluent were collected, while 2 mL fractions were collected during the elution. Fractions containing pure protein were identified by SDS-PAGE gel (BioRad, 3450123). Amicon Ultra 30 kDa MWCO filters (Millipore Sigma, UFC803024) were then used to combine, concentrate, and buffer exchange the purified protein fractions into 10 mM Tris, pH 7.4 (American Bio, AB14044). Protein concentration was determined by NanoDrop One Microvolume UV-Vis Spectrophotometer (Thermo-Fisher) before storage at -80 °C.

**Mucinase digestion**

All proteins were first digested with either SmE, ImpA (NEB, P0761), or OgpA (Genovis, G2-OP1-020), prior to any further processing. All solutions were prepared using MS grade water (Thermo Scientific, 51140). For the structural characterization and mapping of TIM-1 and TIM-4 as well as the analysis of GP1bα, 6-8 µg of protein were used for each digest. All other digests were performed using 2 µg of protein. Each glycoprotein was digested with the O-glycoproteases in a total volume of approximately 15 µL of fresh 50 mM ammonium bicarbonate (AmBic) (Honeywell Fluka, 40867) overnight at 37 °C. Digestions with SmE were conducted at an enzyme to substrate ratio of 1:10, while ImpA and OgpA were digested according to commercial instructions. When sialidase was used, it was added alongside the O-glycoproteases at concentrations in accordance with the manufacturer protocol (NEB, P0720).

**SDS-PAGE analysis**

Fractions from SmE protein purification were run on a 4-12% Criterion XT BisTris gel (Bio-Rad, 3450123) in MES XT buffer (Bio-Rad, 1610789) at 180 V for 60 min, alongside Precision Plus All Blue Protein Standard (Bio-Rad,

1610373). Digested proteins were separated on a 4-12% Criterion XT BisTris gel (Bio-Rad, 3450123) in MOPS XT buffer (Bio-Rad, 1610788) at 180 V for 60 min, alongside Blue Easy Protein Ladder (NIPPON Genetics, MWP06). All protein gels were imaged on a LI-COR Odyssey instrument following a 30 min incubation with Aquastain (Bulldog Bio, AS001000).

**Mass spectrometry sample preparation**

After mucinase digestion, dithiothreitol (DTT) (Sigma Aldrich, D0632) was added to a concentration of 2 mM and allowed to react at 65 °C for 20 min followed by alkylation in 5 mM iodoacetamide (IAA) (Sigma Aldrich, I1149) for 15 min in the dark at room temperature.

For samples where presence of N-glycosylation interfered with identification of potential O-glycosylation sites, a PNGaseF (NEB, P0705) digestion was performed. This included the characterization of C1-INH, while secondary files for TIM1 and TIM3 were generated after gaps in coverage indicated N-glycosylation. The concentrated enzyme was diluted 1:10 and 1 µL of the diluted stock was used for each 2 µg of protein. After allowing the enzyme to react overnight, the protein was buffer exchanged into 50 mM AmBic using 10 kDa MWCO filters (Merck Millipore, UFC501024).

Proteins with fewer sites of glycosylation (i.e. Fetuin, TIM-3, and C1-INH) underwent an additional digestion by adding sequencing-grade trypsin (Promega, V5111) in a 1:50 enzyme:substrate (E:S) ratio for 6 hours at 37 °C. All reactions were quenched by adding 1 µL of formic acid (Thermo Scientific, 85178) and diluted to a volume of 200 µL prior to desalting. Desalting was performed using 10 mg Strata-X 33 µm polymeric reversed phase SPE columns (Phenomenex, 8B-S100-AAK). Each column was activated using 500 µL acetonitrile (ACN) (Honeywell, LC015) followed by of 500 µL 0.1% formic acid, 500 µL 0.1% formic acid in 40% ACN, and equilibration with two additions of 500 µL 0.1% formic acid. After equilibration, the samples were added to the column and rinsed twice with 200 µL 0.1% formic acid. The columns were transferred to a 1.5 mL tube for elution by two additions of 150 µL 0.1% formic acid in 40% ACN. The eluent was then dried using a vacuum concentrator (LabConco) prior to reconstitution in 10 µL of 0.1% formic acid.

**Mass spectrometry data acquisition**

Samples were analyzed by online nanoflow liquid chromatography-tandem mass spectrometry using an Orbitrap Eclipse Tribrid mass spectrometer (Thermo Fisher Scientific) coupled to a Dionex UltiMate 3000 HPLC (Thermo Fisher Scientific). For each analysis, 4 µL was injected onto an Acclaim PepMap 100 column packed with 2 cm of 5 µm C18 material (Thermo Fisher, 164564) using 0.1% formic acid in water (solvent A). Peptides were then separated on a 15 cm PepMap RSLC EASY-Spray C18 column packed with 2 µm C18 material (Thermo Fisher, ES904) using a gradient from 0-35% solvent B (0.1% formic acid with 80% acetonitrile) in 60 min.

Full scan MS1 spectra were collected at a resolution of 60,000, an automatic gain control (AGC) target of 3e5, and a mass range from 300 to 1500 m/z. Dynamic exclusion was enabled with a repeat count of 2, repeat duration of 7 s, and exclusion duration of 7 s. Only charge states 2 to 6 were selected for fragmentation. MS2s were generated at top speed for 3 seconds. Higher-energy collisional dissociation (HCD) was performed on all selected precursor masses with the following parameters: isolation window of 2 m/z, 29% normalized collision energy, orbitrap detection (resolution of 7,500), maximum inject time of 50 ms, and a standard AGC target. An additional electron transfer dissociation (ETD) fragmentation of the same precursor was triggered if 1) the precursor mass was between 300 and 1,500 m/z and 2) 3 of 8 HexNAc or NeuAc fingerprint ions (126.055, 138.055, 144.07, 168.065, 186.076, 204.086, 274.092, and 292.103) were present at ± 0.1 m/z and greater than 5% relative intensity. Two files were collected for each sample: the first collected an ETD scan with supplemental energy (EThcD) while the second method collected a scan without supplemental energy. Both used charge-calibrated ETD reaction times, 100 ms maximum injection time, and standard injection targets. EThcD parameters were as follows: Orbitrap detection (resolution 7,500), calibrated charge-dependent ETD times, 15% nCE for HCD, maximum inject time of 150 ms, and a standard precursor injection target. For the second file, dependent scans were only triggered for precursors below 1000 m/z, and data were collected in the ion trap using a normal scan rate.

**Mass spectrometry data analysis**

Raw files were searched using O-Pair search with MetaMorpheus against directed databases containing the relevant protein sequence.[4] Mass tolerance was set to 10 ppm for MS1's and 20 ppm for MS2's. Met oxidation was set as a variable modification and carbamidomethyl Cys was set as a fixed modification. For samples treated with PNGaseF, Asn deamidation was added as a variable modification. For most samples, we used the default O-glycan database containing 12 common structures. For analysis of GP1bα, the database was based on the previously-determined glycomic profile.[3] Files initially underwent a nonspecific search in order to determine the cleavage specificity of SmE. After the cleavage motif was determined, files generated using only an O-glycoprotease digestion were searched with semi-specific cleavage N-terminal to Ser and Thr and six allowed missed cleavages. Samples treated with trypsin were searched with the same parameters, but also allowed cleavage C-terminal to Arg or Lys. Results were filtered to a q value less than 0.01 and manually validated using Xcalibur software (Thermo Fisher Scientific). Relative abundances were obtained by generating extracted ion chromatograms and determining area under the curve. After abundances were obtained, each file was checked for presence of the identified species. When a peak with matching retention time and mass was present, the peak was validated and the abundance recorded. The mass spectrometry proteomics data have been deposited to the ProteomeXchange Consortium via the PRIDE partner repository with the dataset identifier PXD039583.

**Cell culture**

HeLa cells (ATCC, CCL-2) were grown in T75 flasks (Falcon, 353136) and maintained at 37 °C, 5% $CO_2$. The cells were cultured in DMEM (Gibco, 11965-092) supplemented with 10% fetal bovine serum (FBS, Sigma, F0926), 1% sodium pyruvate (Gibco, 11360-070), and 1% penicillin/streptomycin (Cytiva, SV30010).

**Western blotting for MUC16 on SmE and StcE treated cells**

HeLa cells were seeded in T25 flasks (Falcon, 353109). The following day, the media was removed and dilutions of StcE or SmE (0, 0.05, 0.5, 5, and 50 nM) in Hank's buffered salt solution (HBSS, Gibco, 24020-117) were added for 60 min. Following treatment, media (1 mL) was collected into tubes containing 0.75 µL of 0.5 M EDTA (Invitrogen, 15575-038) to quench the enzymatic reaction. The samples were then concentrated in a 3 kDa spin filter (Millipore, UFC500324). The cells remaining in the flask were washed with an enzyme-free dissociation buffer containing EDTA (Millipore, S-004-C), lifted, and transferred to tubes. The cells were pelleted and washed with PBS (Gibco, 14190-144) twice, then lysed by boiling in 1x NuPAGE LDS Sample Buffer (Invitrogen, NP0008) supplemented with 25 mM DTT at 95 °C for 5 min. Concentrated supernatants were diluted in 4x sample buffer to a final concentration of 1x. Both cell lysates and supernatants were boiled for 5 min at 95 °C and 30 µL of sample was loaded onto a 4-12% Criterion XT BisTris gel (Bio-Rad, 3450124) which was run in MOPS XT buffer (Bio-Rad, 1610788) at 180 V for 90 min. Proteins were transferred to a 0.2 µm nitrocellulose membrane (Bio-Rad, 1620112) using the Trans-Blot Turbo Transfer System (Bio-Rad), at a constant 2.5 A for 15 min. Total protein was quantified using Revert 700 stain (LI-COR Biosciences, 926-11011) before primary antibody (Novus Biologicals, NB600-1468) incubation overnight at 4 °C. An IR800 dye-labeled secondary antibody (LI-COR, 926-32210) was used according to manufacturer's instructions for visualization on a LI-COR Odyssey instrument.

**Cell viability assay**

HeLa cells (CCL-2, ATCC) were seeded in 24-well plates at approximately 20,000 cells per well in 500 µL of media. After 24 hours, SmE and StcE were added at 500, 5, 0.05 and 0 nM. At t = 0, 24, 48, 72, 96 hours post treatment, PrestoBlue (Invitrogen, A13261) was added according to the manufacturer's instructions. After 2 hours, the supernatant was transferred to a black 96 well plate (Thermo Scientific, 237105) for fluorescent readings on a SPECTRAmax GEMINI spectrofluorometer using an excitation wavelength of 544 nm and an emission wavelength of 585 nm. Results were plotted and statistical significance was assessed by two-wayANOVA in GraphPad Prism.

*Computational Methods*

**Molecular Dynamics Simulations**

Protein construction: TIM-3 protein model was built from the following component models: X-ray crystal structure of the TIM-3 IgV domain (PDB 7M41),[5] mucin-domain backbone constructed from the BuildPeptide tool in ROSETTA[6] from the FASTA sequence, and the transmembrane tail and cytoplasmic domains modeled with AlphaFold (AF Q8TDQ0).[7,8] TIM-4 protein model was built from the following component models: X-ray crystal structure of the TIM-4 IgV domain (PDB 5F7H),[9] mucin-domain backbone constructed from the BuildPeptide tool in ROSETTA from the FASTA sequence, and the transmembrane tail and cytoplasmic domains modeled with AlphaFold (AF Q96H15). Each of these domains were then joined together using psfgen in VMDTools.[10]

Glycosylation: An FA2 glycan was chosen for each N-linked glycan positions (list out the N-linked glycan positions) as that was consistent with the known glycoprofile at those positions, and full characterization of N-linked glycan positions is outside the scope of this current work. For all O-linked glycans, the glycan structure at each position with highest population, as determined by MS, was chosen and modeled and constructed onto each site using CHARMM-GUI.[11]

Lipid bilayer insertion: Complete TIM-3 and TIM-4 models were then inserted into lipid bilayer patches with compositions similar to that of mammalian cell membranes (56% POPC, 20% CHL, 11% POPI, 9% POPE and 4% PSM).[12,13]

Solvation and Neutralization: Finally, the TIM-3 and TIM-4 systems were embedded into orthorhombic boxes, explicitly solvated with TIP3 water molecules, and neutralized to a concentration of 150mM of NaCl, resulting in systems of 846,793 and 2,122,863 million atoms, respectively. See **Table S1** for complete system breakdown:

| | TIM-3 | TIM-4 |
|---|---|---|
| Total #Atoms | 846,793 | 2,122,863 |
| #Protein atoms | 4,316 | 5,336 |
| #Glycan atoms | 1,275 | 3,410 |
| #Water atoms | 781,557 | 2,022,120 |
| #Na/#Cl atoms | 830/735 | 2055/1902 |
| Lipid Bilayer Size (Å x Å) | 130 x 130 | 140 x 140 |
| Box Size (Å x Å x Å) | 140.7 x 150.6 x 473.2 | 177.9 x 170.4 x 818. |

**Table S1. Compositional breakdown of each structure simulated in this work.** Lipid bilayer patch size and box size reflect dimensions at t=0 before any simulation was conducted.

Molecular Dynamics (MD) Simulations: All MD simulations were performed with NAMD2.14[14] and CHARMM64m all-atom additive force fields[15–21] on a private supercomputer in the Triton Shared Computing Cluster hosted by the San Diego Supercomputer Center.[22] *Lipid tail minimization and melting:* All atoms except lipid tails were held fixed according to a Lagrangian constraint (i.e. the "fix" command in NAMD), while lipid tails were subjected to 10,000 steps of Steepest Descent minimization. Then, a heating step was performed wherein, with constraints on all atoms except for lipid tails, the system temperature was incrementally increased from 10 K to 310 K for 0.5 ns at 1 fs/step. *Total system minimization and equilibration:* Following lipid tail melting, the Lagrangian constraints were removed from all atoms, but an energetic restraint (1kcal/mol/Å) was applied to all protein and glycan atoms. The complete system was then subjected to 10,000 steps of Steepest Descent minimization, followed by 0.5 ns of equilibration at 310 K (at a 1 fs timestep). After free total minimization and restrained equilibration, the TIM-3 and TIM-4 structures were branched to perform replicas of the following MD simulation protocols. TIM-3 was branched into 4 replicas and TIM-4 was branched into 3 replicas. *Free equilibration:* Finally, all restraints were removed (thus no restraints or constraints on the system at all) and all atoms were allowed to equilibrate for 0.5 ns at 310 K (1 fs/step). *Production:* A total of 700 ns and 830 ns (2 fs/step) were collected for TIM-3 and -4, respectively, see **Table S2** for a breakdown of per replica sampling. MDAnalysis was then used to perform all of the resultant analyses from MD simulations.[23,24]

| | TIM-3 (ns) | TIM-4 (ns) |
|---|---|---|
| Rep 1 | 238.8 | 260.9 |
| Rep 2 | 152.2 | 309.9 |
| Rep 3 | 131.3 | 261.0 |
| Rep 4 (TIM-3 only) | 177.1 | – |
| Total | 699.7 | 831.8 |

**Table S2. Complete breakdown of total simulation time for TIM-3 and TIM-4.**

Persistence Length Calculations: The polymer module in MDAnalysis[23,24] was used to calculate persistence length of TIM-3 and TIM-4 mucin domains by defining a mucin polymer as the N, CA, and C, atoms along the backbone for the following residue selections: TIM-3, residues 133 to 198, TIM-4: residues 137 to 310. Per replica and per TIM protein, persistence length was calculated using the last 1000 frames of simulation. We calculated persistence lengths from each replica trajectory, and thus we have reported the average and standard deviation of persistence lengths calculated for each TIM mucin domain over the three replicas.

End-to-end distance calculations: The distances module in MDAnalysis[23,24] was used to calculate the distance in Angstroms (Å) between the center of mass of the last residue of the globular IgV domain and the center of mass of the first residue of the transmembrane helical domain for every frame in each simulation trajectory. For TIM-3 these first and last residues were selected as 133 and 198, respectively, and for TIM-4 these residues were 137 and 310, respectively. We then normalized the calculated distances by the total number of protein residues within each mucin domain.

Bending Angle Calculations: To calculate the bending angle, we used MDAnalysis[23,24] to select the center of mass of the first, middle, and final protein residue within each mucin domain. We then used these positions to calculate vectors: one from the middle residue to the first residue of each mucin domain, and one from the middle residue and the last residue of each mucin domain. For TIM-3, the first, middle, and last residues were selected as residues 131, 166, and 202, respectively. For TIM-4, the first, middle, and last residues were selected as residues 135, 225, and 314, respectively. We then calculated the angle between these vectors for each frame for all three replica trajectories. We plotted these results as normalized density histograms.

**Mucinase Structure Overlays, Molecular Docking, & TIM-4 Grafting**

Structural comparison and docking were performed using Molecular Operating Environment (MOE) 2020.09. The X-ray crystal structures of ImpA,[25,26] AM0627,[27,28] ZmpB,[25] ZmpC,[29] and BT4244[25] were superimposed with the AlphaFold-predicted structures[7,8] of SmE, AM0908, and AM1514 using the catalytic histidine and glutamate residues to guide alignment.[30]

Following this, a ligand was generated over several steps for use in docking studies. First, the bisglycosylated ligand cocrystallized with AM0627[27] was placed into the analogous location of the superimposed SmE model structure. The amino acid residues of the peptide were then mutated to match a TIM-4 sequence (Ala189-Val190-Phe191-Thr192*-Thr193*-Ala194, where the asterisk indicates glycosylation) that was found to be cleaved by SmE but not ImpA. The peptide's N terminus was acetylated and its C terminus was N-methylated to better approximate the steric/electronic environment of a substrate. While holding the SmE model structure fixed, the glycopeptide was allowed to preliminarily minimize in the Amber10:EHT forcefield[31] using restraints to ensure contacts were formed between (1) the backbones of the glycopeptide and the beta strand of the active site as well as (2) the GalNAc moiety and the side chains of the conserved residues Trp240 and Asn268.

During our glycoproteomic mapping, we observed high occupancy of H1N1A1 (or smaller fragments thereof) modifying Thr192 at P1 and H2N2A1 (or smaller fragments thereof) modifying Thr193 at P1' (**Figure S18**). Similar to previous work,[32,33] the P1 glycan was generated by grafting a sialic acid residue onto the 3-OH of the Gal moiety, using the glycan bound to the GspB siglec domain (PDB 5IUC) as a template.[34] The P1' glycan was similarly generated by grafting the remaining GalNAc, Gal, and Sia residues onto the 6-OH of the GalNAc moiety branching from P1' using the glycan present on PSGL-1 in its cocrystal structure with P-selectin (PDB 1G1S).[35]

Following this, the resulting glycopeptide underwent conformational search, holding all atoms except for the newly-grafted glycans fixed, using LowModeMD to generate a library of over 1300 different conformations of these sugar residues.[36] Each conformer underwent virtual screen, holding the mutant SmED245A enzyme (replacing the catalytic residue to facilitate docking) rigid while allowing the glycopeptide ligand to move freely. The ligand-enzyme complexes with the top 100 docking scores were then used in induced fit docking, keeping both the ligand and the enzyme free. This stepwise search, screen, and docking process allowed us to thoroughly explore conformational space of the ligand while minimizing computational resources.

Following this, the docked ligand was then grafted together with TIM-4 fragments from dynamics simulations to generate larger glycopeptide ligands. First, data from the TIM-4 simulation were analyzed to identify frames containing structures that have Ramachandran angles for Val190 and Phe191 that were within ±15° of those in the docked structure (Val190: $\varphi$ = -140°, $\psi$ = 152°; Phe191: $\varphi$ = -136°, $\psi$ = 102°). Corresponding fragments (Pro175-Phe191) of ten different structures were manually superimposed on the docked structure, using Val190 and Phe191 to guide placement. Finally, the docked ligand (Val190-Ala194) was grafted onto each fragment (Pro175-Ala189) to generate ten larger glycopeptides used to identify potential for interaction with the mucin-binding module of SmE.
